# Disruption of the standard kinetochore in holocentric *Cuscuta* species

**DOI:** 10.1101/2023.01.04.522735

**Authors:** Neumann Pavel, Ludmila Oliveira, Tae-Soo Jang, Petr Novák, Andrea Koblížková, Veit Schubert, Andreas Houben, Jiří Macas

## Abstract

Segregation of chromosomes depends on the centromere. Most species are monocentric, with the centromere restricted to a single region per chromosome. In some organisms, monocentric organization changed to holocentric, in which the centromere activity is distributed over the entire chromosome length. However, the causes and consequences of this transition are poorly understood. Here, we show that the transition in the genus *Cuscuta* was associated with dramatic changes in the kinetochore, a protein complex that mediates the attachment of chromosomes to microtubules. We found that in holocentric *Cuscuta* species the KNL2 genes were lost; the CENP-C, KNL1, and ZWINT1 genes were truncated; the centromeric localization of CENH3, CENP-C, KNL1, MIS12, and NDC80 proteins was disrupted; and the spindle assembly checkpoint (SAC) was degenerated. Our results demonstrate that holocentric *Cuscuta* species lost the ability to form a standard kinetochore and do not employ SAC to control the attachment of microtubules to chromosomes.

## Introduction

Faithful segregation of chromosomes during mitosis and meiosis depends on the centromere, a chromosomal domain that facilitates attachment of chromosomes to spindle microtubules. In monocentric chromosomes, the centromere is localized at a single site per chromosome, which is morphologically discernible as a primary constriction. Holocentric chromosomes, on the other hand, lack this primary constriction and instead have the centromere domains distributed along almost the entire chromosome length. Holocentricity evolved from monocentric organization independently several times during the evolution of both plants and animals ^1^; however, the causes of the transitions are still enigmatic. This is primarily because only a few holocentric species have been studied so far and because most groups of holocentric species evolved from the monocentric ancestors a long time ago, making the factors involved in the transition elusive.

In most species, the centromere is epigenetically determined by the presence of CENH3, a centromere-specific variant of histone H3 that replaces the canonical H3 histones in centromeric nucleosomes ^2^. At the same time, CENH3 serves as the basis for the kinetochore, a complex multiprotein structure that mediates the connection between centromeric chromatin and the microtubules of the mitotic spindle in most species. The backbone of the kinetochore consists of the constitutive centromere associated network (CCAN), which connects the kinetochore with centromeric chromatin, and the KMN network, which constitutes an interface towards spindle microtubules ^3,4^. The function of the kinetochore is regulated by additional proteins, the most studied of which belong to the spindle assembly checkpoint (SAC) ^5,6^ and the chromosome passenger complex (CPC) ^7–9^.

The role of CENH3 in centromere determination predicts that the transition from monocentric to holocentric centromere organization requires the formation of CENH3-containing domains along entire chromosomes. Indeed, in the few holocentric species studied to date, CENH3 is typically localized along the entire poleward surface of each chromatid where microtubules attach ^10,11^. An exception are holocentric insects that lack CENH3 and use an alternative pathway of kinetochore assembly that depends on CENP-T protein ^12–14^.

Recently, we identified the first exception in plants, in *Cuscuta europaea*, which belongs to the holocentric subgenus *Cuscuta* of the parasitic plant genus *Cuscuta* (Convolvulaceae) ^15^. In this species, the chromosomes restrict CENH3 to only one to three heterochromatin bands, despite being attached to the mitotic spindle along their entire length. This suggests that CENH3 has either lost its centromere function in this species or acts in parallel with an additional CENH3-independent mechanism of kinetochore assembly. Since monocentric relatives of *C. europaea* from the sister subgenus, *Grammica*, and the more distant subgenus, *Monogynella*, have CENH3 localized specifically in primary constrictions ^16^, it is plausible that the peculiar CENH3 localization in *C. europaea* resulted from changes in kinetochore assembly that were linked to the transition to holocentricity in the subgenus *Cuscuta*. However, how kinetochore assembly has changed and whether these changes are related to the transition to holocentricity remains unknown.

In this study, we addressed these questions by comparing the repertoire of major structural and regulatory kinetochore proteins and their chromosomal localization between two *Cuscuta* species from the holocentric subgenus *Cuscuta* (*C. europaea* and *C. epithymum*), two monocentric *Cuscuta* species from the sister subgenus *Grammica* (*C. australis* and *C. campestris*), and *Ipomoea nil*, which was included as an outgroup Convolvulaceae species. To obtain high-quality data for gene identification in the two holocentric *Cuscuta* species, we sequenced both their genomes and transcriptomes. The chromosomal localization of kinetochore proteins was determined using antibodies developed against key proteins representing different subcomponents of the kinetochore. Comparison of the results between monocentric and holocentric species allowed us to uncover an unprecedented level of changes that occurred specifically in the holocentric species and thus likely played an important role in the transition to holocentricity in *Cuscuta*.

## Results

### Transition to holocentricity in *Cuscuta* was associated with massive changes of kinetochore protein genes

Sequencing of the holocentric species *C. europaea* and *C. epithymum* resulted in genome assemblies of 975.8 Mb (N50 = 17.9 Mb) and 997 Mb (N50 = 3.3 Mb), respectively (Supplementary Note 1, and Supplementary Table 1). The completeness of gene space and quality of gene prediction were assessed using BUSCO and were comparable to genome assemblies previously published for the monocentric *Cuscuta* relatives *C. australis* and *C. campestris* (Supplementary Fig. 1). The quality of gene prediction in the genome assembly was also verified by the independent assembly of the transcriptomes, which showed similar results following BUSCO analysis (Supplementary Table 2). To identify kinetochore protein sequences in the species selected for this study, we created a sequence database of 29 structural and regulatory kinetochore proteins known in plants. First, we used the database as a query for blastp searches to identify homologous protein sequences in the monocentric species *C. australis, C. campestris*, and *Ipomoea nil*. The identified sequences were manually verified and corrected when needed, and added to the database to improve its sensitivity for homologous protein recognition. The improved database was then used for blastp searches in the two holocentric *Cuscuta* species. Comparison of the identified kinetochore protein genes revealed that all 29 tested genes are present and mostly intact in the monocentric species, whereas in the holocentric species some of the genes are either absent, significantly truncated, or duplicated accompanied by a higher rate of sequence divergence (Fig. 1a, Supplementary Table 3 and Supplementary Data 1).

**Fig. 1.**
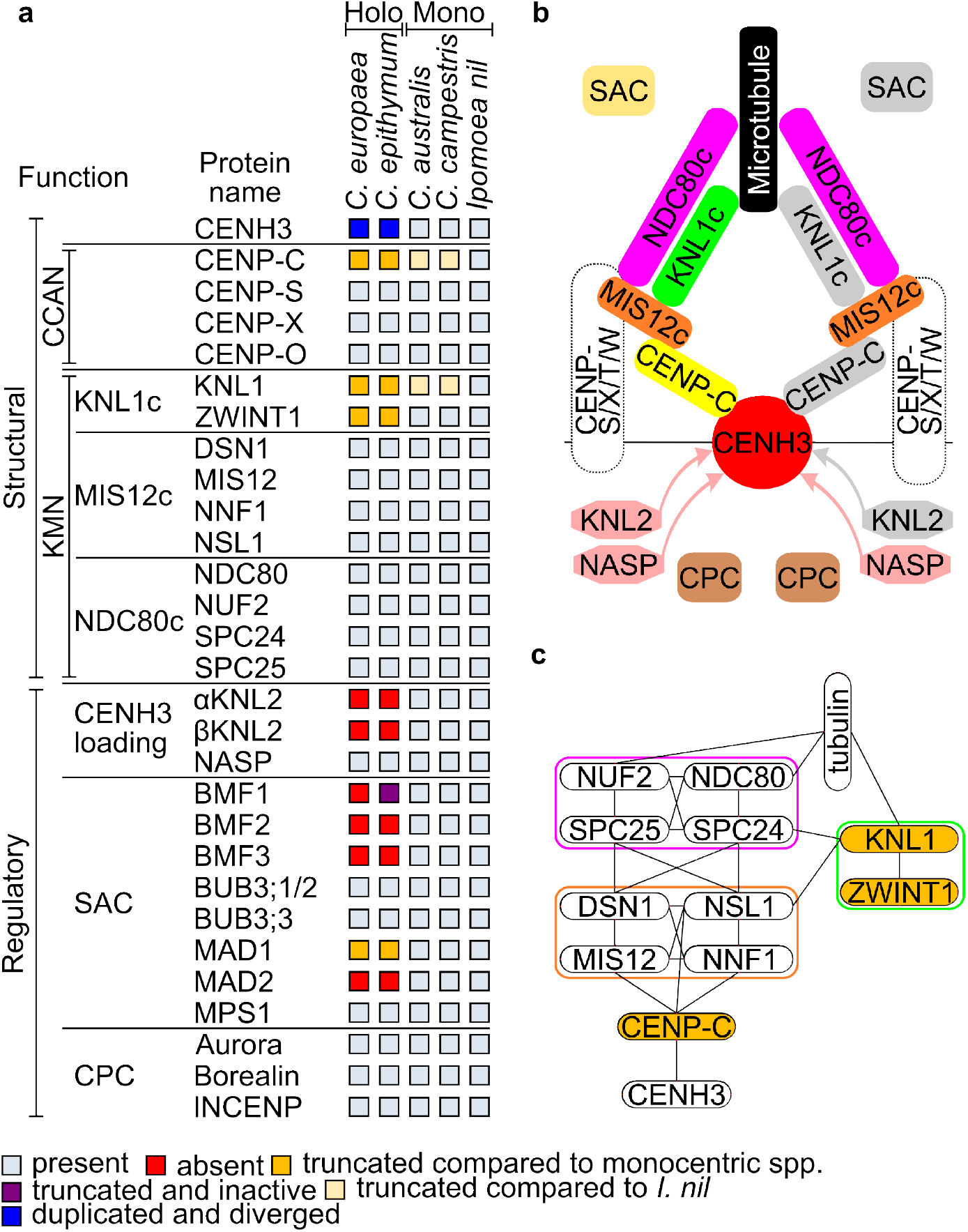
The repertoire of structural and regulatory kinetochore proteins analyzed in this study. **a**, Results of the survey of the protein sequences. **b**, Simplified schematic illustration of kinetochore structure ^3,4^. Proteins or complexes containing proteins that have been truncated or lost in holocentric *Cuscuta* species are highlighted in gray on the right. Centromeric chromatin is determined by the presence of CENH3. The deposition of CENH3 in plants depends on the KNL2 and NASP proteins. The outer kinetochore consists of the KMN network, which includes three subcomplexes, KNL1c, MIS12c, and NDC80c. The connection between centromeric chromatin and the KMN network is mediated by CENP-C. Some metazoan species have an alternative pathway of kinetochore assembly based on CENP-T. CENP-T forms a complex with CENP-S, CENP-X, and CENP-W, and also interacts with NDC80c and MIS12c ^12,59^. Because the plant homologs of CENP-T are not known, it is not clear whether the CENP-T pathway also exists in plants. The precise spatiotemporal and orderly progression of mitosis is ensured by the activity of regulatory kinetochore proteins belonging to the spindle assembly checkpoint (SAC) and the chromosome passenger complex (CPC). The SAC monitors the state of chromosome attachment to spindle microtubules and prevents the transition from metaphase to anaphase until all sister chromatids are attached to microtubules ^6^. The CPC is involved in mitotic checkpoint activity, destabilizes improperly attached spindle microtubules, and promotes axial shortening of chromosome arms during anaphase ^7–9^. **c**, Schematic illustration of the interactions between the proteins forming the CENP-C pathway of kinetochore assembly. Proteins truncated in holocentric *Cuscuta* species are highlighted in orange. The interactions were drawn based on findings in yeast and humans ^24,60–65^ but likely also occur in plants.

The lost genes included both eudicotyledonous plant homologs of *KNL2*, referred to as *αKNL2* and *βKNL2* ^17^, and four of eight spindle assembly checkpoint (SAC) genes, namely, *BMF1, BMF2, BMF3*, and *MAD2* (Fig. 1a). Their absence was in all cases confirmed by comparison of genomic loci possessing these genes in *C. australis* with the orthologous loci in *C. europaea* and *C. epithymum* (Supplementary Figs. 2 and 3), as well as by their absence in genome-independent transcriptome assemblies. The only exception was *BMF1* whose transcriptionally inactive fragment still remains in *C. epithymum* (Supplementary Fig. 3). Large gene truncations took place in three structural kinetochore protein genes, including *CENP-C, KNL1*, and *ZWINT1*, and the SAC gene *MAD1* (Figs. 2 and 3 and Supplementary Figs. 4 and 5). Finally, the CENH3 gene in holocentric species was found to have duplicated once in the common ancestor of *C. europaea* and *C. epithymum*, and once independently in each of the two species. The diversification of the duplicated CENH3 genes in holocentric species resulted in considerably higher protein sequence variability for CENH3 compared with monocentric *Grammica* species, suggesting that they evolved more rapidly (Supplementary Figs. 6, 7, 8 and 9).

**Fig. 2.**
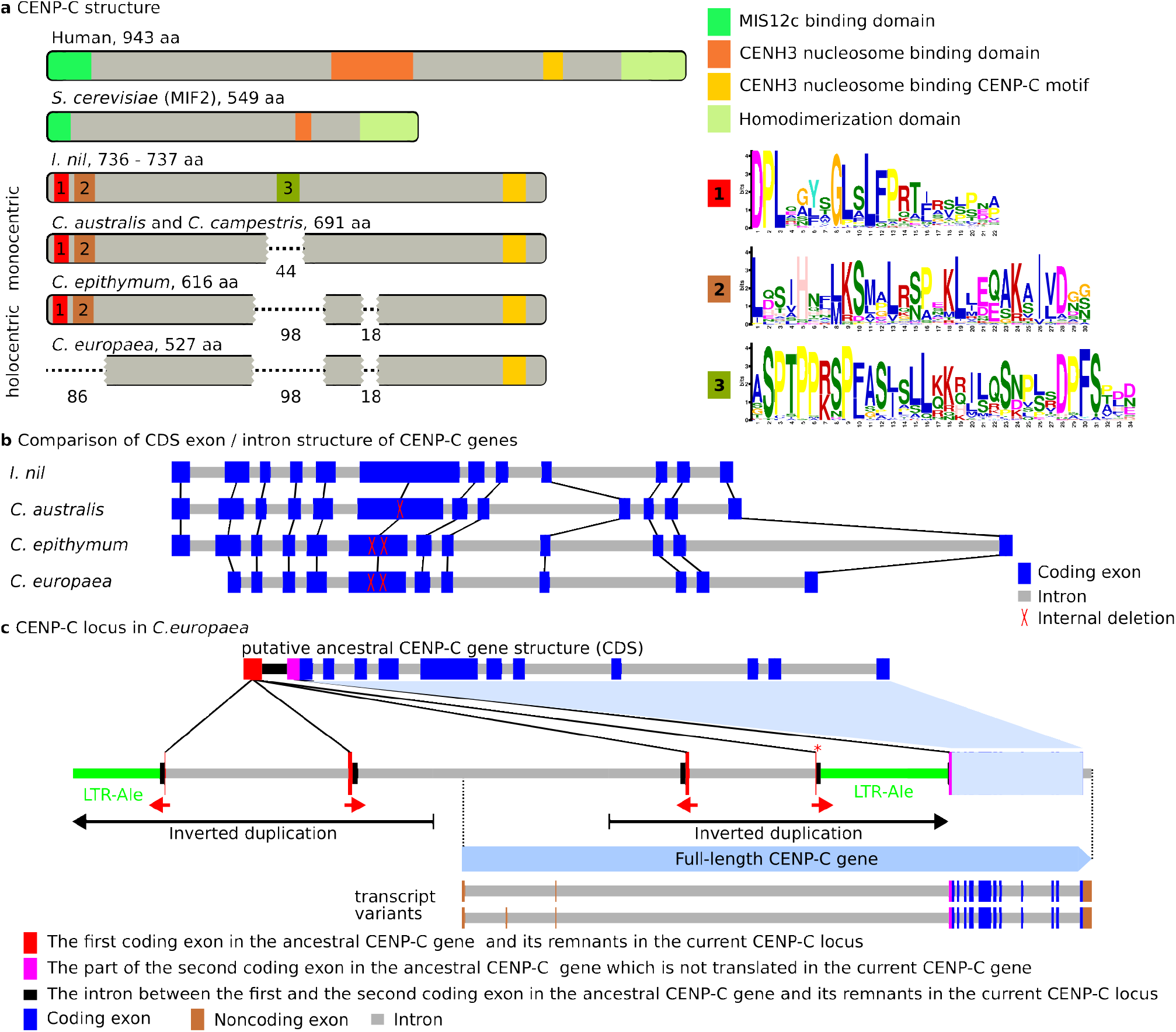
Truncation of CENP-C in *Cuscuta* species. **a**, Comparison of domain structure between human, S*accharomyces cerevisiae*, and monocentric and holocentric Convolvulaceae species. Human and yeast CENP-C sequences are divergent, but the positions of the functional domains are conserved ^65,66^. Compared with *I. nil*, CENP-C is truncated in both monocentric and holocentric *Cuscuta* species, but with more extensive truncations in the latter species (the missing parts are shown as dashed lines, and the numbers below indicate their length). The N-terminal truncation in *C. europaea* and the internal truncations in all *Cuscuta* species resulted in the loss of domains recognized by MEME as conserved in dicotyledonous plants, indicating their functional importance. The sequence logos of these domains are shown on the right. **b**, Comparison of CDS exon-intron structure of CENP-C genes. The exon-intron structure is conserved in all Convolvulaceae species (the orthologous exons are connected with black lines). The internal truncations of CENP-C proteins in *Cuscuta* species are due to deletions in the sixth coding exon. **c**, Schematic illustration of the CENP-C gene locus in *C. europaea*. The current CENP-C locus is compared with the putative ancestral CENP-C gene structure (top), which was reconstructed by adding the missing region from *C. epithymum*. The original CENP-C gene gradually changed by a short inverted duplication of the first coding exon and part of the following intron (red arrows), a partial deletion in the first coding exon that remained in the correct orientation (marked with a red asterisk), the insertion of the Ty1/Copia LTR retrotransposon Ale (green), and a large inverted duplication (black arrows). The remnants of the first ancestral coding exon became part of the intron. The second ancestral coding exon was retained and became the first coding exon of the gene in present-day *C. europaea*.

**Fig. 3.**
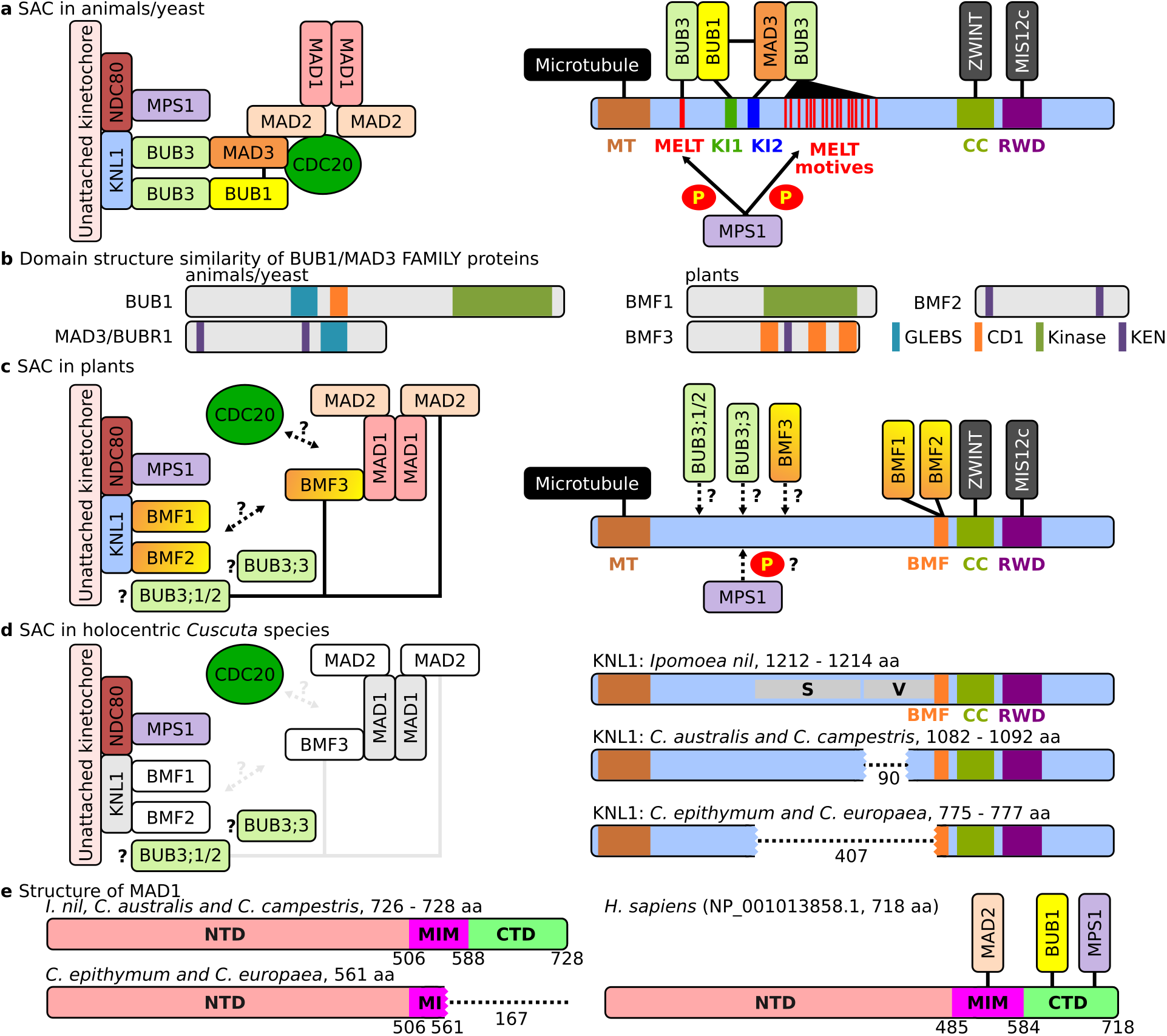
Schematic illustration of interactions between SAC and KNL1 in animals/yeasts, plants, and holocentric *Cuscuta* species. Schematics were adapted from ^6,47^, with modifications to reflect the results of other studies cited below. **a**, SAC and KNL1 in animals and yeast. (Left) SAC is activated on kinetochores that are not attached to microtubules, and its formation is initiated by MPS1, a kinase that phosphorylates MELT repeats in KNL1. Phosphorylated KNL1 serves as a binding platform for SAC, which interacts with CDC20 to form the mitotic checkpoint complex, preventing entry into anaphase. (Right) Schematic representation of the protein binding domains in KNL1 (drawn after ^67^). Protein interactions in both schematics are shown as adjacent rectangles or black lines. **b**, Domain organization in BUB1/MAD3 family (BMF) proteins in animals/yeasts and their plant counterparts BMF1-BMF3 (drawn after ^47^). **c**, SAC and KNL1 in plants. (Left) The architecture of the plant SAC differs from that in animals/yeasts ^47,48,68,69^, and the function and interactions of some SAC proteins are not yet known (dashed lines with question marks). (Right) Plant KNL1 lacks MELT, KI1, and KI2 domains and binds BMF1 and BMF2 proteins via the BMF domain near the C-terminus ^48^. **d**, SAC and KNL1 in holocentric *Cuscuta* species. (Left) SAC is severely impaired by the absence or truncation of several proteins (white and gray boxes, respectively). (Right) Truncation of KNL1 in *Cuscuta* species as compared to *I. nil*. The region missing in monocentric *Cuscuta* species corresponds to a highly variable region (V), whereas the region missing in the holocentric species also includes a segment that shares sequence similarity to KNL1 from various plant species (S). The truncations are depicted as dotted lines, and their lengths are indicated by the numbers below. **e**, Structure of MAD1. N-terminal domain (NTD), MAD2 interaction motif (MIM), and C-terminal domain (CTD) were determined by comparison with human MAD2 ^70^. C-terminal truncation of MAD2 in holocentric *Cuscuta* species resulted in the loss of domains interacting with MAD2, BUB1, and MPS1 in humans.

Given the function of proteins that are either missing or truncated, the changes are likely to have had a substantial impact on kinetochore assembly and function at multiple levels, from CENH3 loading (absence of KNL2) and kinetochore assembly (truncation of CENP-C, KNL1, and ZWINT1), to regulation of its function (absence of several key proteins of SAC) (Fig. 1b,c).

### CENH3 histones do not have holocentric-like distribution in holocentric Cuscuta species

Since KNL2 is essential for proper loading of CENH3 to centromeres ^17–20^, the loss of both *αKNL2* and *βKNL2* in holocentric *Cuscuta* species is likely to have a serious impact on CENH3 localization. On holocentric chromosomes, CENH3 is expected to specifically localize along the poleward side of each chromatid. In contrast to this expectation, we have previously shown that CENH3 occurs in all but one prominent transversal heterochromatin band in *C. europaea* and that CENH3 distribution does not correlate with the distribution of mitotic spindle attachment sites detected with antibodies against α-tubulin (^15^ and Fig. 4a,b). To determine the localization of CENH3 in *C. epithymum*, we developed three antibodies against different N-terminal sequence variants of the proteins. Although the antibodies were made to recognize all CENH3 protein sequence variants present in the tested plant, none of them produced a signal on chromosomes and nuclei that could be distinguished from the background (Supplementary Fig. 10a-g). On the other hand, two of the antibodies developed for *C. epithymum* detected CENH3 in the heterochromatin domains in *C. europaea* (Supplementary Fig. 10c,e), demonstrating that they were functional for *in situ* detection. These results suggest that CENH3 is either not present in chromatin in *C. epithymum* or that its levels are considerably lower than in *C. europaea*, and thus below the limits of detection for the applied *in situ* immunodetection technique. Despite the absence of CENH3 signal, α-tubulin immunostaining revealed attachment of mitotic spindle microtubules to chromosomes along their poleward sides, confirming the holocentric nature of chromosomes in *C. epithymum* (Fig. 4c). This was in contrast to monocentric *Cuscuta* spp., which had microtubules attached only to CENH3 containing domains (Fig. 4d and data not shown). These results suggest that CENH3 does not function as a foundational kinetochore protein in holocentric *Cuscuta* species.

**Fig. 4.**
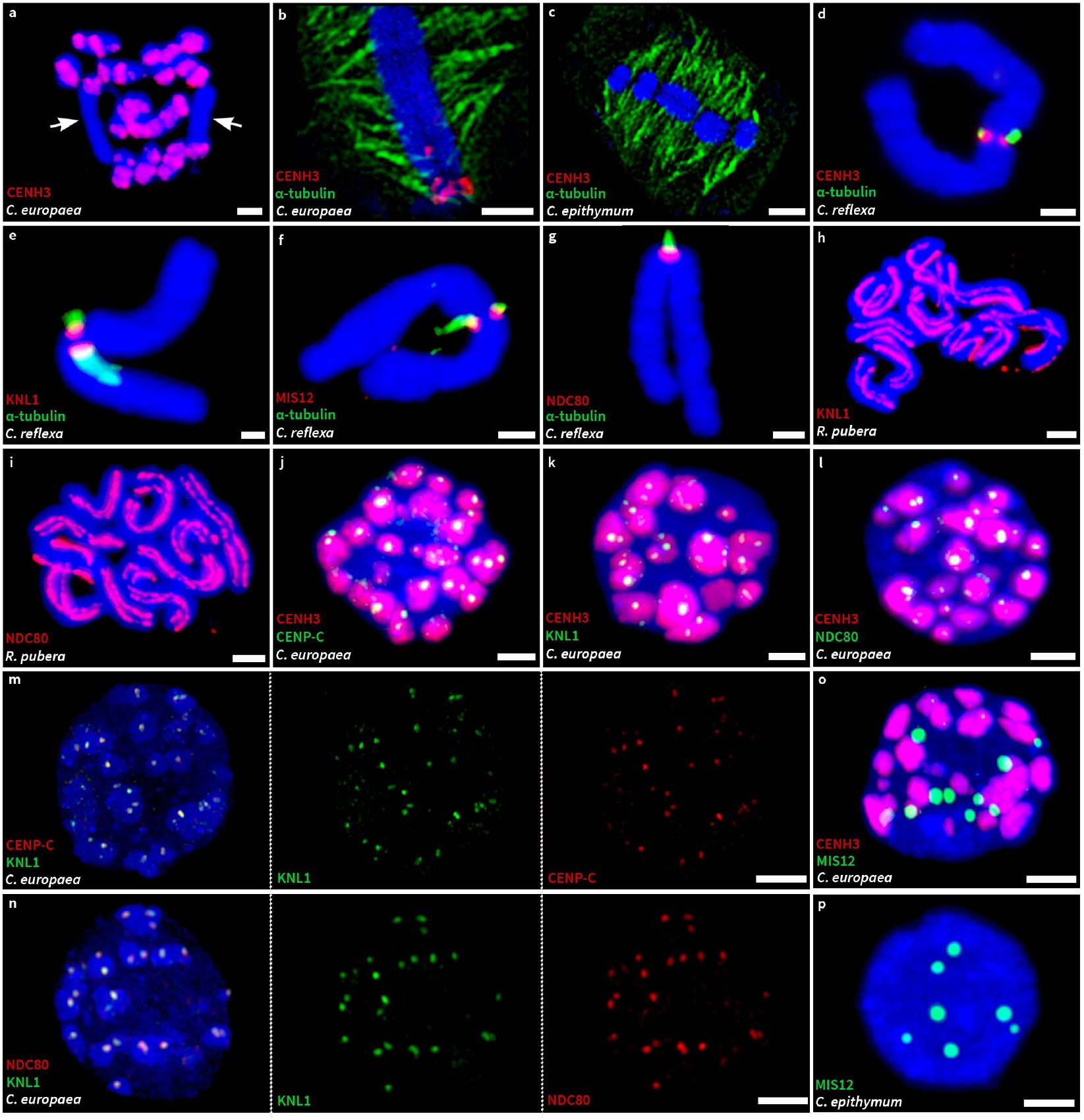
*In situ* immunodetection of structural kinetochore proteins and α-tubulin. **a**, Detection of CENH3 on mitotic chromosomes in *C. europaea*. Arrows indicate chromosomes 1, which possess a single subtelomeric CENH3-containing domain, while the majority of the chromosome lacks CENH3 signals. **b**, Detection of CENH3 and α-tubulin on selected chromosome 1 in *C. europaea*. The image is a single optical section selected from an 3D-SIM image stack showing that microtubules of the mitotic spindle are evenly attached to the chromosome at its poleward sides and along its entire length, independent of the occurrence of CENH3 signals. **c**, Detection of CENH3 and α-tubulin in *C. epithymum*. The image is a single optical section selected from an 3D-SIM image stack showing even distribution of microtubules of the mitotic spindle despite the absence of CENH3 signals. **d-g**, Detection of α-tubulin with either CENH3 (d), KNL1 (e), MIS12 (f), and NDC80 (g) on selected *C. reflexa* chromosomes. All four proteins are specifically localized on the surface of primary constriction where microtubules attach. **h-i**, Detection of KNL1 (h) and NDC80 (i) in *Rhynchospora pubera*. Both proteins show holocentromere-characteristic distribution of both proteins along the entire length of all chromosomes. **j-l**, Detection of CENH3 with either CENP-C (j), KNL1 (k), or NDC80 (l) in interphase nuclei of *C. europaea*. CENP-C, KNL1, and NDC80 are localized in small domains embedded in much larger CENH3-containing heterochromatin domains. The images were reconstructed using maximum-intensity projection from 3D-SIM image stacks. **m-n**, Detection of KNL1 with either CENP-C (m) or NDC80 (n), showing that all three proteins are colocalized. The images were reconstructed using maximum-intensity projection from 3D-SIM image stacks. **o**, Detection of MIS12 and CENH3 in an interphase nucleus of *C. europaea*, showing that the two proteins are not colocalized. **p**, Detection of MIS12 in interphase nucleus of *C. epithymum*. The spatial visualizations of nuclei shown in k, l, o, and p are available as Supplementary Movies 1-4. Chromosomes were stained with DAPI (blue). Scale bars = 2 µm.

### Kinetochore assembly is impaired in holocentric Cuscuta species

The chromosomal distribution of CENH3 together with the truncation of three structural kinetochore proteins suggested that kinetochore assembly may be impaired in holocentric *Cuscuta* species. To test whether the kinetochore assembles along the poleward chromosome surface, as expected for holocentric chromosomes, we examined the localization of CENP-C, which is a linker between CENH3 and the KMN network, and of MIS12, KNL1, and NDC80, which represent the three complexes of the KMN network (Fig. 1b). Antibodies were developed against peptides designed from domains that were conserved in the holocentric species. However, owing to high sequence similarity between species, it was likely that the antibodies against KNL1, NDC80, and MIS12 would also recognize homologous proteins from monocentric *Cuscuta* species. Indeed, when these antibodies were used for *in situ* detection, monocentromeres in *C. australis* as well as in *C. reflexa* from the more distant subgenus *Monogynella* were labeled, demonstrating the functionality of the antibodies (Fig. 4e-g and Supplementary Fig. 11). The antibodies against KNL1 and NDC80 proved to be particularly versatile, functioning even in *Rhynchospora pubera*, an evolutionarily very distant plant species with holocentric chromosomes, where they detected holocentromere-characteristic signals for both proteins (Fig. 4h,i). In agreement with the lack of CENH3 signal in *C. epithymum*, CENP-C, KNL1 and NDC80 were not detected on either mitotic chromosomes or in interphase nuclei in this species (data not shown). In *C. europaea*, these three proteins were detected in small subdomains embedded within CENH3-containing heterochromatin during interphase but not on mitotic chromosomes (Fig. 4j-l, Supplementary Movie 1, and data not shown). Simultaneous *in situ* detection of KNL1 with either CENP-C or NDC80 revealed that these proteins fully colocalized (Fig. 4m,n and Supplementary Movies 2 and 3). These results suggest that the assembly of the kinetochore during interphase in *C. europaea* still depends, at least in part, on the presence of CENH3, but that kinetochore organization is disrupted before cells enter mitosis. Strikingly, MIS12 was detected in 2 - 16 (n = 100) discrete nuclear domains during interphase in both holocentric species (Fig. 4o,p). In *C. europaea*, these domains were always located away from the CENH3-containing heterochromatin (Fig. 4o and Supplementary Movie 4), indicating that MIS12 has become independent of CENP-C and the KMN network proteins.

### *Conventional SAC is abolished in holocentric* Cuscuta *species*

To test if the regulatory kinetochore complexes form on chromosomes in holocentric *Cuscuta* species despite the absence of the tested kinetochore proteins and the massive loss of the SAC genes observed, we raised antibodies against BUB3;1/2 and Borealin, which are components of the SAC and CPC, respectively. While the BUB3;1/2 antibodies produced monocentric-like signals on chromosomes in *C. australis* and *C. reflexa*, and holocentromere-like signals in *Rhynchospora pubera*, BUB3;1/2 was not detectable on chromosomes in holocentric *Cuscuta* species (Fig. 5a-c and data not shown). On the other hand, the antibodies against Borealin labeled the chromosomes in the region around areas of sister chromatid cohesion at centromeres in monocentric *C. reflexa* and along the entire chromosome length in both holocentric *Cuscuta* species (Fig. 5d-f). These results indicate that the conventional SAC is abolished, while the CPC maintains at least some of its functions in holocentric *Cuscuta* species.

**Fig. 5.**
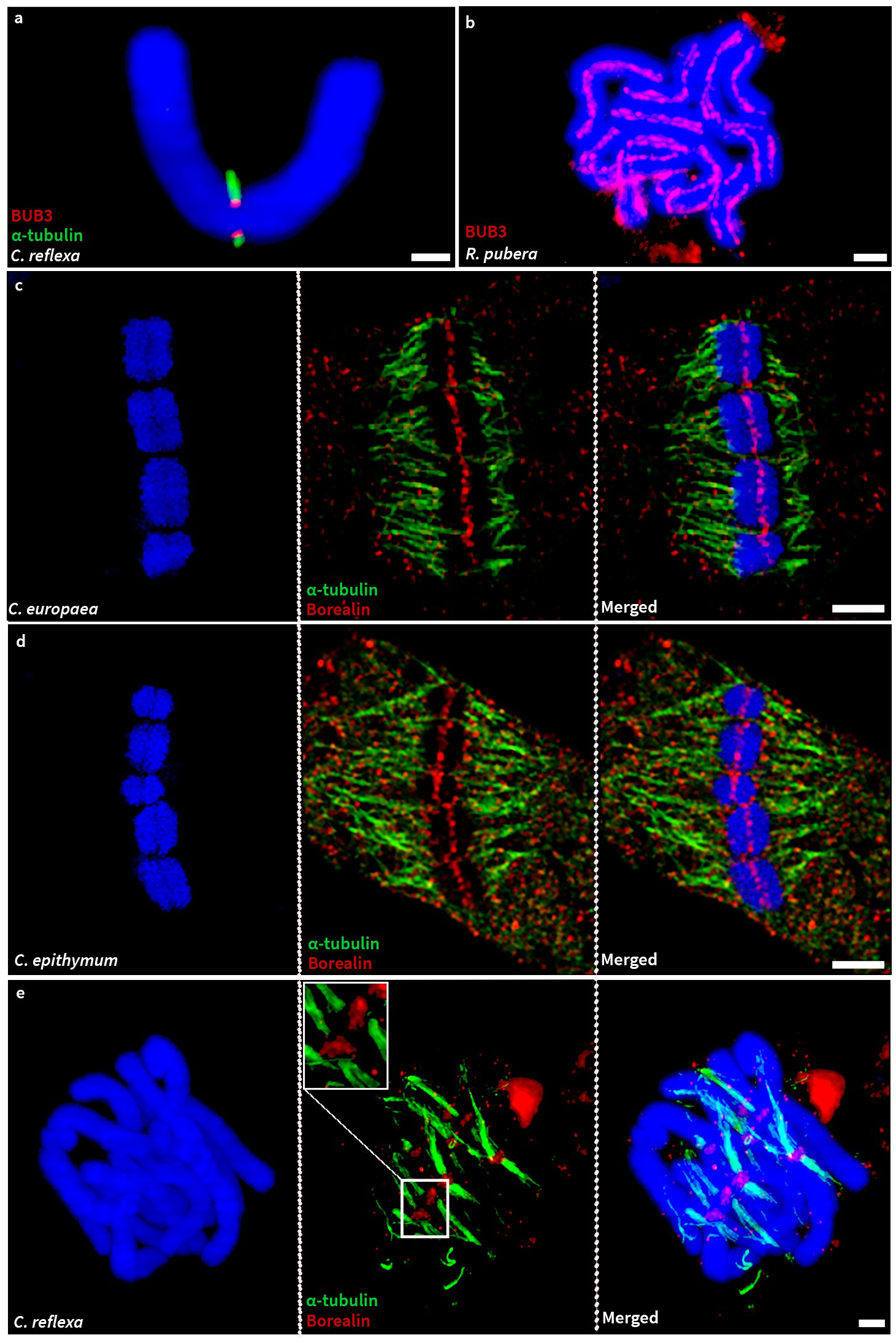
*In situ* immunodetection of BUB3;1/2 and Borealin. **a**, Simultaneous detection of BUB3;1/2 and α-tubulin on mitotic chromosomes in *C. reflexa*. The image shows that BUB3;1/2 is specifically localized on the surface of the primary constriction where microtubules attach. **b**, Detection of BUB3;1/2 on mitotic chromosomes in *R. pubera*, showing holocentromere-characteristic distribution of the signals along the entire length of all chromosomes. **c-d**, Simultaneous detection of Borealin and α-tubulin on mitotic chromosomes in *C. europaea* (c) and *C. epithymum* (d). The images show single optical slices selected from 3D-SIM image stacks. **e**, Simultaneous detection of Borealin and α-tubulin on mitotic chromosomes in *C. reflexa*. Chromosomes were stained with DAPI (blue). Scale bars = 2 µm.

## Discussion

The peculiar CENH3 localization in *C. europaea* described in our previous study ^15^ suggested that the transition to holocentricity in the genus *Cuscuta* may have been associated with the formation of a CENH3-independent kinetochore assembly. In this study, we have demonstrated that the transition to holocentricity in *Cuscuta* species was associated with extensive changes in structural and regulatory kinetochore protein genes, and disruption of both standard kinetochore assembly and SAC regulation of mitotic chromosome segregation. This distinguishes holocentric *Cuscuta* species from both the holocentric nematode *Caenorhabditis elegans*, which use the CENH3-CENP-C pathway of kinetochore assembly ^21^, and holocentric insects, in which the CENH3-CENP-C pathway of kinetochore assembly was lost and replaced by the CENP-T pathway ^12–14^ (Fig. 6).

**Fig. 6.**
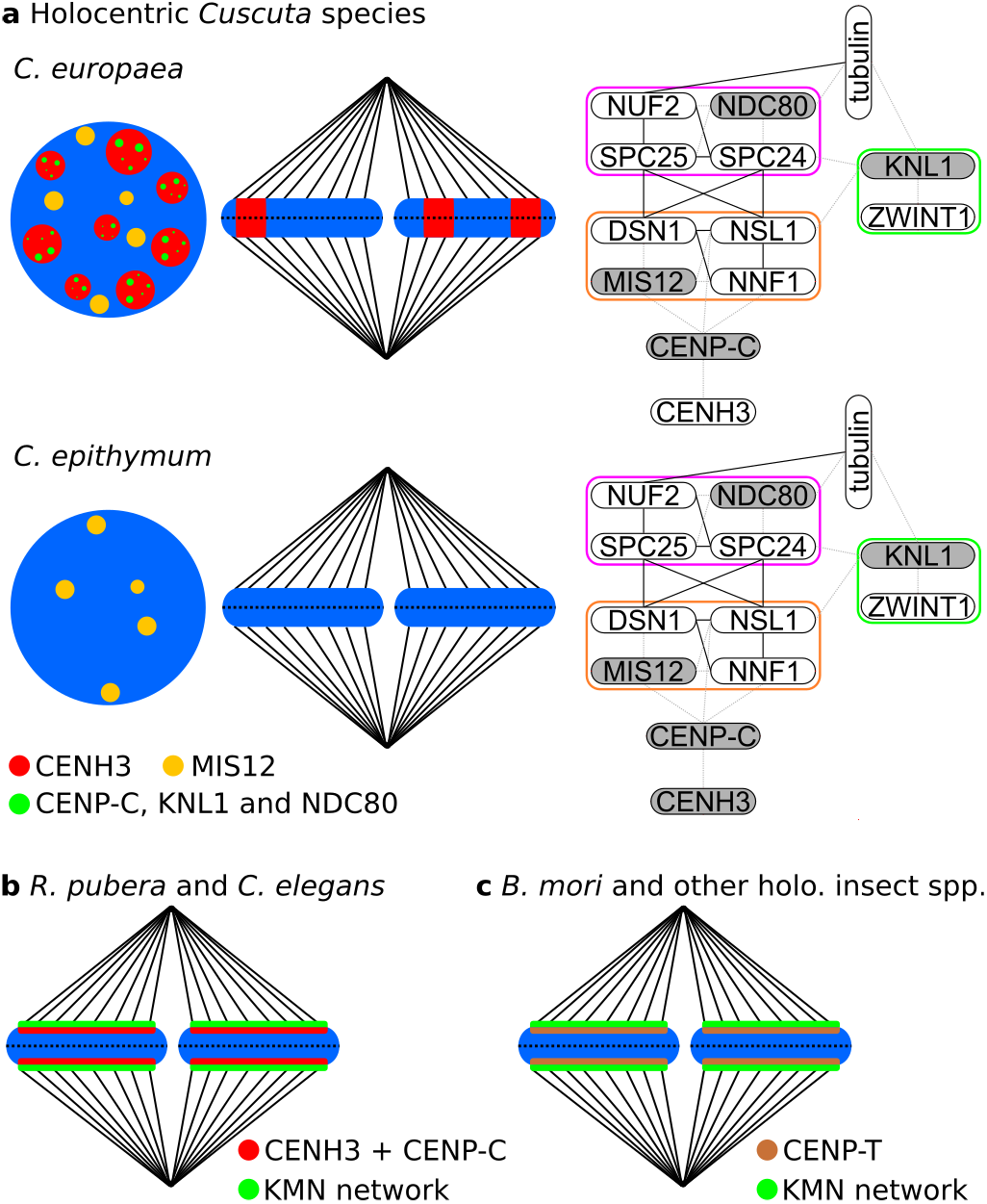
Comparison of kinetochore structure between holocentric *Cuscuta* species and other previously studied holocentric species. **a**, (Left) Summary of the distribution of structural kinetochore proteins examined in this study in *C. europaea* and *C. epithymum*. In both *Cuscuta* species, the microtubules of the mitotic spindle are attached to the chromosomes along their entire length, indicating their holocentric nature. In *C. europaea*, CENH3 is specifically localized in transverse heterochromatin bands rather than on the poleward surface along the entire chromosome length. During interphase, the kinetochore proteins CENP-C, KNL1, and NDC80 are colocalized in small areas within CENH3-containing heterochromatin, whereas MIS12 occurs at separate, discrete sites. None of these proteins were detected on mitotic chromosomes. In *C. epithymum*, which lacks conspicuous heterochromatin domains, CENH3, CENP-C, KNL1, and NDC80 were not detected in interphase nuclei or on mitotic chromosomes, whereas MIS12 was detected at several discrete sites in interphase nuclei. (Right) Schematic illustrations of the interactions between the proteins forming the CENP-C pathway of kinetochore assembly from Fig. 1c, where the proteins that were examined but not detected on mitotic chromosomes are shaded in gray and the resulting missing interactions are shown as gray dashed lines. They show that the absence of these proteins likely disrupts overall kinetochore assembly. **b**, Kinetochore formation on holocentromeres in *R. pubera* and *Caenorhabditis elegans*. Centromere domains are determined by the presence of CENH3. On mitotic chromosomes, they form a continuous layer on the poleward surface of each chromatid where the kinetochore forms and spindle microtubules attach. The KMN network of the outer kinetochore is connected to the CENH3-containing nucleosomes via the CENP-C protein. **c**, Kinetochore formation on holocentromeres in *Bombyx mori* and other holocentric insect species. These species lack CENH3 and the KMN network is linked to chromosomes via the CENP-T protein.

We hypothesize that one of the most important changes in the evolution of holocentric *Cuscuta* species was the loss of KNL2. In *C. elegans*, RNAi depletion of KNL2 leads to a reduction in the presence of CENH3 to levels undetectable by immunodetection, resulting in chromosome segregation defects and embryonic lethality ^18,19^. Similar phenotypes have been observed in KNL2 mutants in other species, including *A. thaliana*, demonstrating the general importance of KNL2 for CENH3 loading ^17,20^. Therefore, the depletion/absence of CENH3 in *C. epithymum* chromatin could be due to the absence of both KNL2 variants. On the other hand, it is puzzling that CENH3 accumulates in heterochromatin domains in *C. europaea* despite the loss of KNL2. Given that all heterochromatin domains that contain CENH3 possess the same repetitive sequences, whereas the heterochromatin domain that lacks these repeats also lacks CENH3 ^15,22^, the incorporation of CENH3 into these domains could be DNA sequence-dependent. In light of the importance of KNL2 and CENH3 for centromere determination and kinetochore assembly, it is surprising that the loss of KNL2 in both holocentric *Cuscuta* species, the depletion/absence of CENH3 on chromosomes in *C. epithymum*, and the peculiar CENH3 distribution on chromosomes in *C. europaea* are neither lethal nor cause chromosome segregation defects. The simplest explanation is that CENH3 is no longer necessary for correct chromosome segregation in holocentric *Cuscuta* species (Supplementary Fig. 12).

The absence of detectable levels of structural kinetochore proteins on mitotic chromosomes in holocentric *Cuscuta* species is in contrast not only to monocentric *Cuscuta* species but also to the holocentric-like distribution of NDC80 and KNL1 in *R. pubera* (Cyperaceae), which was used as a holocentric control plant in this study (Fig. 4). This suggests that the formation of the standard kinetochore is disrupted in holocentric *Cuscuta* species. In *C. epithymum*, this could be primarily a direct consequence of the depletion/absence of CENH3 on the chromosomes. In *C. europaea*, the causes of kinetochore disruption must be different because CENH3-containing heterochromatin is present throughout the cell cycle and partially colocalizes with CENP-C, KNL1, and NDC80 proteins during interphase. The reasons why the putative complex of kinetochore proteins formed during interphase disappears at the onset of mitosis are not clear. Considering that three structural kinetochore proteins are truncated (Fig. 1a), one possibility is that the complex falls apart because of disrupted interactions between kinetochore components (Fig. 1c). The truncation of CENP-C may be the most critical because CENP-C is the only protein known to link centromeric chromatin to the outer kinetochore in plants (Figs. 1b,c and 2a). Although the N-terminus of CENP-C is divergent in sequence between eukaryotes, it has been shown to bind MIS12c in both humans and yeast, indicating a conserved function ^23–25^. Given that this function is also conserved in plants, the N-terminal truncation of CENP-C in *C. europaea* should interfere with MIS12c binding. Consistent with this notion, we found that MIS12 does not colocalize with CENP-C and accumulates in discrete domains that are clearly separated from CENH3-containing domains (Fig. 4l and Supplementary Movie 4). While the colocalization of CENP-C, KNL1, and NDC80 suggests that the kinetochore assembles during interphase, despite the absence of MIS12, the complex may not be sufficiently stable to survive mitosis. The N-terminal truncation of CENP-C is, however, unlikely to cause the disappearance of the protein itself because the N-terminus is not required for the binding of centromeric nucleosomes (Fig. 2a and ^26^). Although the internal portion of CENP-C contains a domain that binds centromeric nucleosomes in humans and yeast (Fig. 2a), the high sequence divergence of CENP-C prevented us from determining by a sequence similarity-based approach whether it overlaps with the region lost in *C. europaea*. On the other hand, the large size disparity between the domains containing CENH3 and CENP-C (Fig. 4j,o and Supplementary Movies 1 and 2) suggests that there is an imbalance between the levels of the two proteins that may reflect inefficient binding of CENP-C to CENH3.

The results discussed above support a model in which holocentric *Cuscuta* species either use substantially reduced kinetochores lacking CENH3, CENP-C, KNL1, MIS12, and NDC80 or, more likely, have evolved a completely novel mechanism of chromosome attachment to the mitotic spindle. This conclusion is also supported by the degeneracy of SAC genes that would have been required had the kinetochore been present and functioning in a conventional manner. Alternative kinetochores have already been described in Kinetoplastida, most of which have lost CENH3 and all CCAN and KMN genes. They consist of proteins that probably evolved from meiotic components of chromosome synapsis and homologous recombination machinery ^27,28^. Moreover, kinetochore-independent chromosomal movement along the spindle, facilitated by kinesin motor proteins, has been described for acentric chromosomes in *Drosophila* neuroblasts ^29,30^ and for chromatin knobs in maize ^31,32^.

Overall, we have shown that the transition to holocentricity in *Cuscuta* species was unique among all species studied to date. It was accompanied, and perhaps even triggered, by the degeneration of standard kinetochore structure and regulation and the formation of a novel mechanism for chromosome attachment to microtubules. The insights gained in this study provide the basis for future studies aimed at uncovering the plasticity of kinetochore assembly and discovering as yet unknown principles of chromosome segregation.

## Material and Methods

### Plant material

Seeds of *C. europaea* (serial number: 0101147) were obtained from the Royal Botanic Garden (Ardingly, UK). *C. epithymum* plants were collected from a natural population at “U Cáby” (Kroclov, Czech Republic). Seeds of *C. australis* and *C. campestris* were provided by Prof. Jianqiang Wu (Kunming Institute of Botany, Chinese Academy of Sciences, Kunming, China) and Dr. Chnar Fathoulla (University of Salahaddin, Kurdistan Region, Iraq), respectively. *C. reflexa* Roxb. plant was obtained from the Botanic Gardens of the Rhenish Friedrich-Wilhelm University (Bonn, Germany). *Cuscuta* plants were cultivated on the following host plant species: *Urtica dioica* (*C. europaea*), *Betonica officinalis* and *Coleus blumei* (*C. epithymum*), *Ocimum basilicum* (*C. australis* and *C. campestris*), or *Pelargonium zonale* (*C. reflexa*). Plants of *R. pubera* were obtained from Dr. André Marques (Max Planck Institute for Plant Breeding Research, Cologne, Germany).

### Genome sequencing and assembly

DNA for Illumina and Pac-Bio sequencing was isolated using the CTAB method from nuclei extracted from young shoots of *C. europaea* and *C. epithymum* as described previously ^33^. Shotgun Illumina paired-end sequencing of DNA was performed by the Brigham Young University (Provo, UT, USA) and Admera Health (South Plainfield, NJ, USA). High molecular weight nuclear DNA used for Oxford nanopore sequencing was isolated using a modified CTAB protocol as described previously ^34^. Nanopore sequencing was performed as described ^22^. Detailed information about all genome sequence datasets produced in this study is provided in Supplementary Table 4.

Illumina paired-end reads and Oxford nanopore reads were assembled using MaSuRCA ^35^. PacBio HiFi reads were assembled using Hifiasm assembler (v0.15.5-r350; ^36^) with default parameters for PacBio HiFi sequence reads. Since the quality of the HiFi-based assemblies were considerably better than those generated by MaSuRCA (Supplementary Table 1), they were selected for submission to European Nucleotide Archive (https://www.ebi.ac.uk/ena/browser/home; Accession numbers: ERZ12293622 (*C. europaea*) and ERZ12293623 (*C. epithymum*)). Completeness and contiguity of assemblies were evaluated using BUSCO (v5.2.2; ^37^) and QUAST (v5.0.2; ^38^). Genome characteristics were evaluated using kmer analysis and the jellyfish program ^39^ with kmer length 21 and 51 for Illumina and PacBio HIFI sequence reads, respectively. Heterozygosity was estimated using GenomeScope program ^40^.

### Transcriptome sequencing, assembly and gene prediction

Total RNA was isolated using the Trizol method. Preliminary sequencing for de-novo transcriptome assemblies of *C. epithymum, C. europaea*, and *C. campestris* was performed at GATC Biotech (Konstanz, Germany) using Illumina technology producing 50bp paired-end reads. In each species, RNA was isolated from shoots and inflorescences, mixed in a 1:1 ratio, treated with DNase I (Ambion, Austin, TX, USA), and then enriched for poly-A fraction using the Dynabeads mRNA purification kit (Thermo Fisher Scientific, Waltham, MA, USA). Deep transcriptome sequencing of *C. epithymum, C. europaea*, and *C. australis* was done using RNA isolated from shoot tips, shoot internodia, or inflorescences at various stages of development. For each species and tissue, the RNA samples were produced in three biological replicates (samples from different plants collected at different time). Subtraction of poly-A RNA using NEBNext Ultra II with a Poly-A Selection kit (New England Biolabs, Ipswich, MA, USA) and poly-A RNA sequencing were performed at Admera Health (South Plainfield, NJ, USA). The sequencing generated more than 500 million 151 nt long paired-end reads for each RNA sample, giving a total yield of about 5 billion reads per species (Supplementary Table 5).

Transcriptomes were de-novo assembled using the Trinity program ^41^ with default options from pair-end reads. Sequences from individual replicates and tissue samples of each species were concatenated before their assembly. The presence of single copy orthologs in the transcriptomes was evaluated using the BUSCO (v.5.2.2) program ^37^. To create gene models, pair-end RNA-Seq Illumina reads were aligned to genome assembly using the STAR program (v2.7.7a; ^42^) with parameters --outSAMstrandField intronMotif --outSAMtype BAM SortedByCoordinate -- alignIntronMax 20000. Each sample was aligned independently. Resulting alignments were merged into a single BAM file using samtools ^43^. Whole length transcripts and genes were then reconstructed using the Stringtie program (v2.1.7; ^44^) with parameters -c 2 -f 0.05. Candidate coding regions within transcript sequences were identified using TransDecoder program (https://github.com/TransDecoder/TransDecoder) with default settings.

Predicted protein sequences from *C. europaea* and *C. epithymum* were compared with published proteomes of *C. campestris, C. australis*, and *Ipomoea nil* using program OrthoFinder (v2.5.2; ^45^) to identify orthologs and orthogroups. Genome assemblies and associated files containing detailed information about predicted gene models, protein and CDS sequences were downloaded from http://plabipd.de/portal/cuscuta-campestris (*C. campestris*) or GenBank (https://www.ncbi.nlm.nih.gov/genbank/; *C. australis*: GCA_003260385.1; *I. nil*: GCF_001879475.1). RNA-seq data for these species were downloaded from the Sequence Read Archive (SRA; https://www.ncbi.nlm.nih.gov/sra) from the following accession numbers: SRR6664647 – SRR6664654 (*C. australis*), ERR1916345 – ERR1916364 (*C. campestris*), and DRR024544 – DRR024549 (*I. nil*). The RNA-seq data produced in this study or downloaded from other studies were used to verify and correct automatically predicted gene models if needed. Manual verification and editing of gene models were performed using Apollo Genome Annotation Editor ^46^.

### Identification and characterization of kinetochore proteins

Structural and regulatory kinetochore protein sequences identified in *A. thaliana* were downloaded from uniprot database and from published studies ^47–49^. These sequences were used for blastp searches to identify their homologs in genome assemblies of *C. australis* and *C. campestris* ^50,51^, representing monocentric *Cuscuta* species, and in *I. nil* ^52^, selected as a monocentric nonparasitic genus of the family Convolvulaceae. All sequences with significant similarity hits were manually inspected to remove false positives, correct erroneous protein sequences, or add additional variants due to alternative splicing. Protein sequences from *A. thaliana* and the three Convolvulaceae species were combined into a reference data set that was used for blastp and tblastn searches to find homologous kinetochore protein genes in holocentric *C. epithymum* and *C. europaea*. The searches were primarily performed in gene and protein sequences predicted using StringTie in the assembly produced from Pac-Bio reads, but the results were verified using the data from the parallel genome assemblies that were made from Illumina and nanopore reads as well as the transcriptome assemblies produced using Trinity.

CENH3 sequences from additional *Cuscuta* species or other plants of the same *Cuscuta* species were obtained from our previous study (*C. campestris, C. japonica* ^15^), identified in transcriptome shotgun assemblies (*C. reflexa, C. campestris*) or other available genome assembly (*C. epithymum*), amplified from RNA using RT-PCR or RACE methods (*C. epithymum*), or reconstructed from available next generation genome sequence data using GRABb and GeneWise programs (*C. americana, C. californica, C. pentagona*; ^53,54^). More detailed information about sources of the CENH3 sequences is provided in Supplementary Table 6.

Sequence alignments were performed using MUSCLE ^55^. Time trees were inferred using ITS and *rbc*L sequences and methods described in our previous study ^16^. ITS and *rbc*L sequences from *C. australis* were reconstructed from Illumina paired end reads (SRA run accession number: SRR5851367) using RepeatExplorer ^56^. A search for conserved sequence motives was performed using MEME ^57^. Sequence logos were generated using WebLogo ^58^. The sources of CENP-C and ZWINT1 sequences used for MEME and WebLogo analyses are provided in the Supplementary Table 7.

### Antibodies

Antibodies to all kinetochore proteins used in this study were custom-produced by GenScript (Piscataway, NJ, USA) or Biomatik (Cambridge, ON, Canada) against peptides designed from regions that were most conserved among *Cuscuta* species and *I. nil*. The particular peptide sequences used for immunization in rabbits were always designed from *C. europaea* kinetochore protein sequences, with the exception of CENH3, which was designed from variable N-termini. The peptide sequences are provided in Supplementary Table 8. Antibody specificity was confirmed using *in situ* immunodetection to identify signals in the primary constrictions of monocentric *Cuscuta* species. The mouse monoclonal antibody to α-tubulin was purchased from Sigma-Aldrich (St. Louis, MO, USA; catalog number: T6199).

Reactivity of the antibodies raised against CENH3 with individual CENH3 variants in *C. europaea* and *C. epithymum* was tested using western blot. Full-length CENH3-coding sequences were cloned into pEXP5-NT/TOPO vector (Invitrogen, Carlsbad, CA, USA) in frame with the N-terminal 6xHis tag-coding sequence. Recombinant proteins were produced in BL21-AI strain of *E. coli* (Invitrogen, Carlsbad, CA, USA) upon induction with isopropyl β-D-thiogalactoside (IPTG). Total protein was extracted using 1× SDS-PAGE buffer according to the manufacturer’s instructions supplied with the pEXP5-NT/TOPO vector, separated on 12% SDS-PAGE gel, and then transferred onto Immobilon-P membrane (Sigma-Aldrich, St. Louis, MO, USA) using TE77XP semi-dry transfer unit (Hoefer, Holliston, MA, USA). Membranes were blocked using 5% skim milk powder in 1× PBS (PBS-M) overnight at 4°C and then incubated for 2 hours at RT with the primary antibody diluted in 1× PBS-M to 2–3 μg/ml. Following six washes in 1× PBS for 10 min at RT each, the antibodies were detected using goat anti-rabbit IgG StarBright Blue 520 secondary antibodies (Bio-Rad, Hercules, CA, USA; catalog number: 12005870) in 1× PBS-M for 1 h at RT. Fluorescent signals were visualized using the Chemidoc MP imaging system (Bio-Rad, Hercules, CA, USA). The presence of recombinant CENH3 proteins on the membrane was always verified by detection with the HisG epitope tag antibody (Thermo Fisher Scientific, Waltham, MA, USA; catalog number: R940-25) and secondary antibody StarBright Blue 700 Goat Anti-Mouse IgG (Bio-Rad, Hercules, CA, USA; catalog number: 12004159).

### *In situ* immunodetection of kinetochore proteins

The biological material (shoot tips for *Cuscuta* and root tips for *Rhynchospora*) was fixed in TRIS-fix buffer (4% formaldehyde, 10 mM Tris, 10 mM Na_2_EDTA, 100 mM NaCl, pH 7.5) for 30 min at 10°C. Infiltration of the fixative was enhanced by applying a vacuum during the first 5 minutes. After fixation, the material was washed in TRIS buffer (10 mM Tris, 10 mM Na 2 EDTA, 100 mM NaCl, pH 7.5) on ice for 30 minutes. For the preparation of chromosomes and nuclei in *Cuscuta* species, the squashing technique was first used after digesting the shoot apical meristems for one hour at 27.4°C in 2% cellulase ONOZUKA R10 (SERVA Electrophoresis, Heidelberg, Germany) and 2% pectinase (MP Biomedicals, Santa Ana, CA, USA). The squashes were performed in either 1× phosphate-buffered saline (PBS) or LB01 (15 mM Tris(hydroxymethyl)aminomethane, 2 mM Na_2_EDTA, 0.5 mM spermine, 80 mM KCI, 20 mM NaCl, 15 mM mercaptoethanol, and 0.1% (v/v) Triton X-100, pH 7.5). With this technique, it was possible to obtain reasonable results, but to minimize background, chromosomes and nuclei were later isolated in suspension as described below. Shoot apical meristems were cut up in 1 ml of cold LB01 using a mechanical homogenizer (Ultra-turrax T8, IKA Z404519). The suspension was filtered through a 48 μm nylon mesh and spun onto slides using a Hettich centrifuge with cytospin chambers. In *Rhynchospora pubera*, formaldehyde-fixed root tip meristems were digested with 2% cellulase ONOZUKA R10 (SERVA Electrophoresis, Heidelberg, Germany) and 2% pectinase (MP Biomedicals, Santa Ana, CA, USA) for one hour at 37 °C. After washing with cold distilled water, meristems were squashed in 1× PBS. Before immunostaining, slides were incubated for 30 minutes at room temperature (RT) in 1× PBS-T1 buffer (1× PBS and 0.5% Triton, pH 7.4) (RT) to increase permeabilization. Slides were washed twice in 1× PBS for 5 minutes at RT and once in 1× PBS-T2 (1× PBS, 0.1% Tween 20, pH 7.4) for 5 minutes at RT. For immunostaining, slides were incubated with primary antibody diluted in 1× PBS-T2 overnight at 4°C. The dilution ratios were as follows: 1:1000 for antibodies to kinetochore proteins and 1:100 for antibodies to α-tubulin (Sigma-Aldrich, St. Louis, MO; catalog number T6199). After washing twice for 5 minutes in 1× PBS at RT, slides were incubated for one hour at RT with the secondary antibody in 1× PBS and then washed twice for 5 minutes in 1× PBS at RT. Primary rabbit and mouse antibodies were detected with goat anti-rabbit Rhodamine Red X (dilution 1:500; Jackson ImmunoResearch, Suffolk, UK; catalog number: 111-295-144) and goat anti-mouse Alexa Fluor 488 (dilution 1:500; Jackson ImmunoResearch; catalog number: 115-545-166), respectively. To distinguish specific signals from background signals caused by nonspecific binding of the secondary antibody, negative control slides were used and subjected to the same treatments as for standard detection, except that the primary antibody was not added. For simultaneous detection of different proteins with two rabbit antibodies, antibodies were labeled directly using Alexa Fluor 488 and Alexa Fluor 568 antibody labeling kits (Thermo Fisher Scientific, Waltham, MA, USA; catalog numbers: A20181 and A20184, respectively) according to the manufacturer’s recommendations. The degree of labeling was determined using a spectrophotometer DS-11 (DeNovix, Wilmington, DE, USA). Before embedding the slides in Vectashield mounting medium (Vector Laboratories, Burlingame, CA) supplemented with 49,6-diamino-2-phenylindole (DAPI), the slides were fixed with 4% formaldehyde in 1× PBS for 10 minutes at RT and then washed twice for 5 minutes in 1× PBS at RT.

### Microscopy

For conventional wide-field fluorescence microscopy, a Zeiss AxioImager.Z2 microscope equipped with an Axiocam 506 mono camera was used along with an Apotome2.0 device for better resolution in the z-axis, which was needed when the images were composed of multiple optical sections. Images were generated using the ZEN 3.2 software (Carl Zeiss GmbH). To capture signals at the super-resolution level (∼120 nm using a 488 nm laser), spatial structured illumination microscopy (3D-SIM) was performed using a 63×/1.4 Oil Plan-Apochromat objective on an Elyra PS.1 microscope system, controlled by the ZENBlack software (Carl Zeiss GmbH). Images were captured using the 405, 488, and 561 nm laser lines for excitation and the appropriate emission filters ^59^. Three-dimensional movies were produced from 3D-SIM image stacks using the Imaris 9.7 (Bitplane) software.

## Acknowledgements

This research was financially supported by grants from the Czech Science Foundation (20-25440S) and the Czech Academy of Sciences (RVO:60077344). Computational resources and data-storage facilities were provided by the ELIXIR-CZ Research Infrastructure Project (LM2018131). We thank to J. Látalová and V. Tetourová for their technical assistance.

## Supplementary Information

### Supplementary Notes

**Supplementary Note 1: Genome Assembly and gene prediction in holocentric *Cuscuta* spp**.

To assemble genome sequences of *C. epithymum* and *C. europaea*, we sequenced the genomic DNA using Illumina, Oxford nanopore, and Pac-Bio Hi-Fi sequencing technologies. Sequence reads from the two former technologies were assembled using MaSuRCA ^1^, whereas Pac-Bio Hi-Fi reads were assembled using Hifiasm ^2^. The latter type of the assembly was considerably better in both species (Supplementary Table 1). The total assembly size in *C. epithymum* was 975 Mbp, which is 1.8-fold bigger than the estimated genome size (1C = 533 Mb) ^3^. This disparity was attributed to high heterozygosity in the sequenced clone, resulting in the presence of two haplotypes in the assembly (Supplementary Table 1). The *C. europaea* genome assembly was 997 Mbp in size, corresponding to about 85% of previously estimated genome (1C = 1,169 Mb). This difference was likely due to the presence of highly abundant satellite DNA repeats, which make up 18% of the genome and are generally difficult to assemble ^3^. Gene prediction using the Stringtie program resulted in 89,521 and 49,635 gene models for *C. epithymum* and *C. europea*, respectively. The almost two-fold higher number of gene models in *C. epithymum* was caused by the presence of two haplotypes in the assembly and thus two alleles for most genes. BUSCO analysis revealed a high proportion of missing genes in both *C. epithymum* and *C. europaea*, but comparison with *C. campestris, C. australis*, and *I. nil* showed that it was not due to poor genome assemblies and/or gene prediction but to a large gene loss that preceded the divergence of monocentric and holocentric *Cuscuta* species (Supplementary Fig. 1). This was also confirmed by BUSCO analysis of the assembly-independent *de novo* transcriptome assemblies (Supplementary Table 2).

### Supplementary Figures

**Supplementary Fig. 1.**
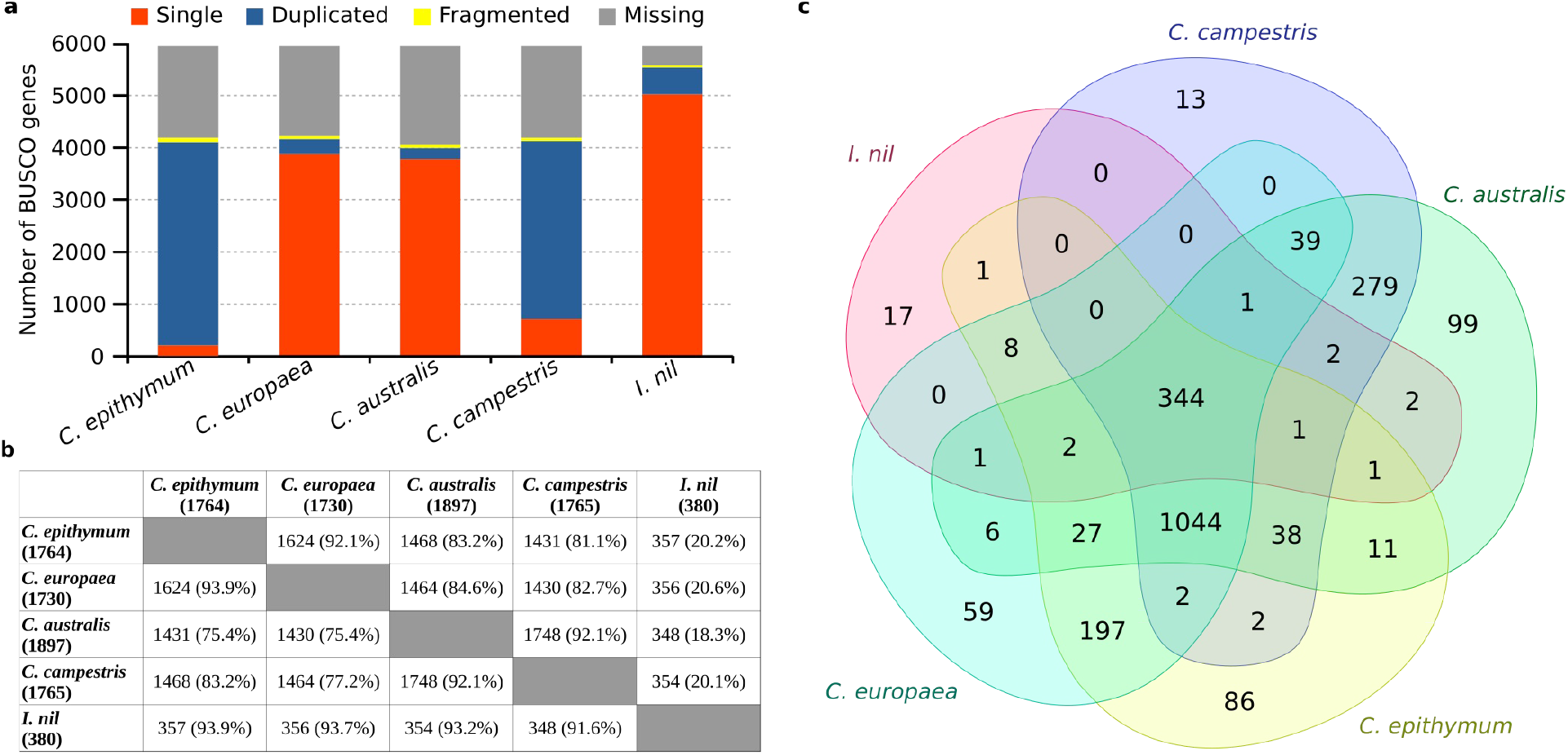
Assessment of the completeness of the gene content in genome assemblies of *C. europaea* and *C. epithymum*. The analysis was done with BUSCO using Solanales_odb10 dataset containing 5590 genes and the results were compared with those obtained for previously published genome assemblies of *C. australis, C. campestris*, and *I. nil* ^4–6^. **a**, Summary of BUSCO results. The number of missing BUSCO genes is similar between *C. europaea* and *C. epithymum* sequenced in this study and monocentric *Cuscuta* species sequenced previously. The high number of duplicated genes in *C. epithymum* and *C. campestris* reflects the presence of two haplotypes and tetraploid origin, respectively. **b**, Pairwise species comparison of missing BUSCO genes. The analysis shows that not only the two holocentric but also the two monocentric species share a high proportion of missing BUSCO genes. The percentages of genes missing for each species shown in the rows are indicated in brackets. **c**, Venn diagram showing overlaps of missing BUSCO genes between all five species. Overall, 344 genes were probably lost before the divergence of the five Convolucaeae species and an additional 1044 genes were lost before divergence of the four *Cuscuta* species. On the other hand, only 86 (1.4%) and 59 (1.0%) BUSCO genes were missing specifically in *C. epithymum* and *C. europaea*, respectively. These results demonstrate that the high number of missing BUSCO genes is not due to poor genome assemblies and/or gene prediction in *C. europaea* and *C. epithymum*, but to relatively massive gene loss that preceded the divergence of monocentric and holocentric *Cuscuta* species.

**Supplementary Fig. 2.**
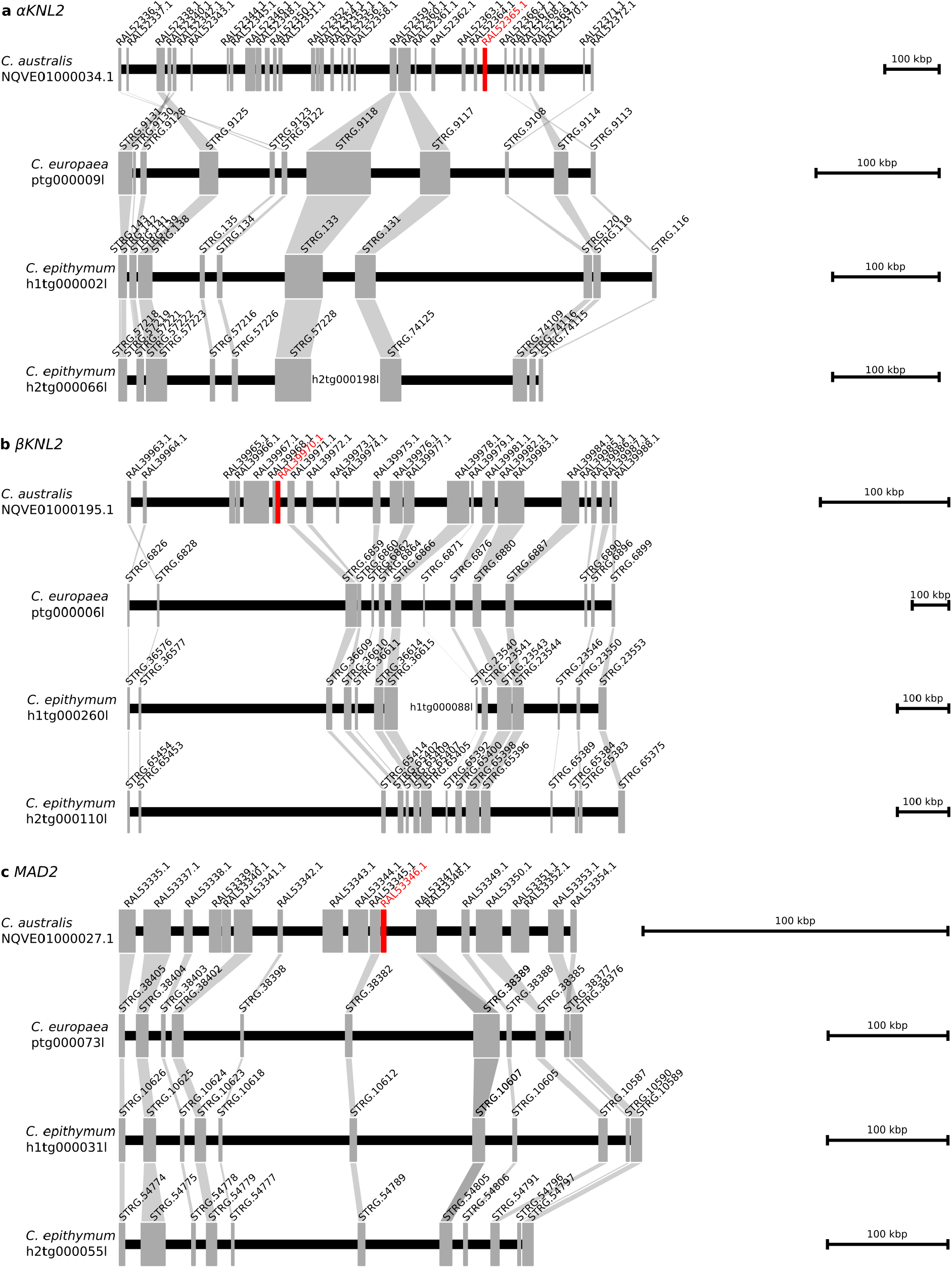
Comparison of orthologous loci, part 1. Comparison of loci possessing *αKNL2, βKNL2*, and *MAD2* genes (highlighted in red) in *C. australis* with orthologous loci in *C. europaea* and *C. epithymum*. As *C. epithymum* has two haplotypes each locus is represented by two contigs.

**Supplementary Fig. 3.**
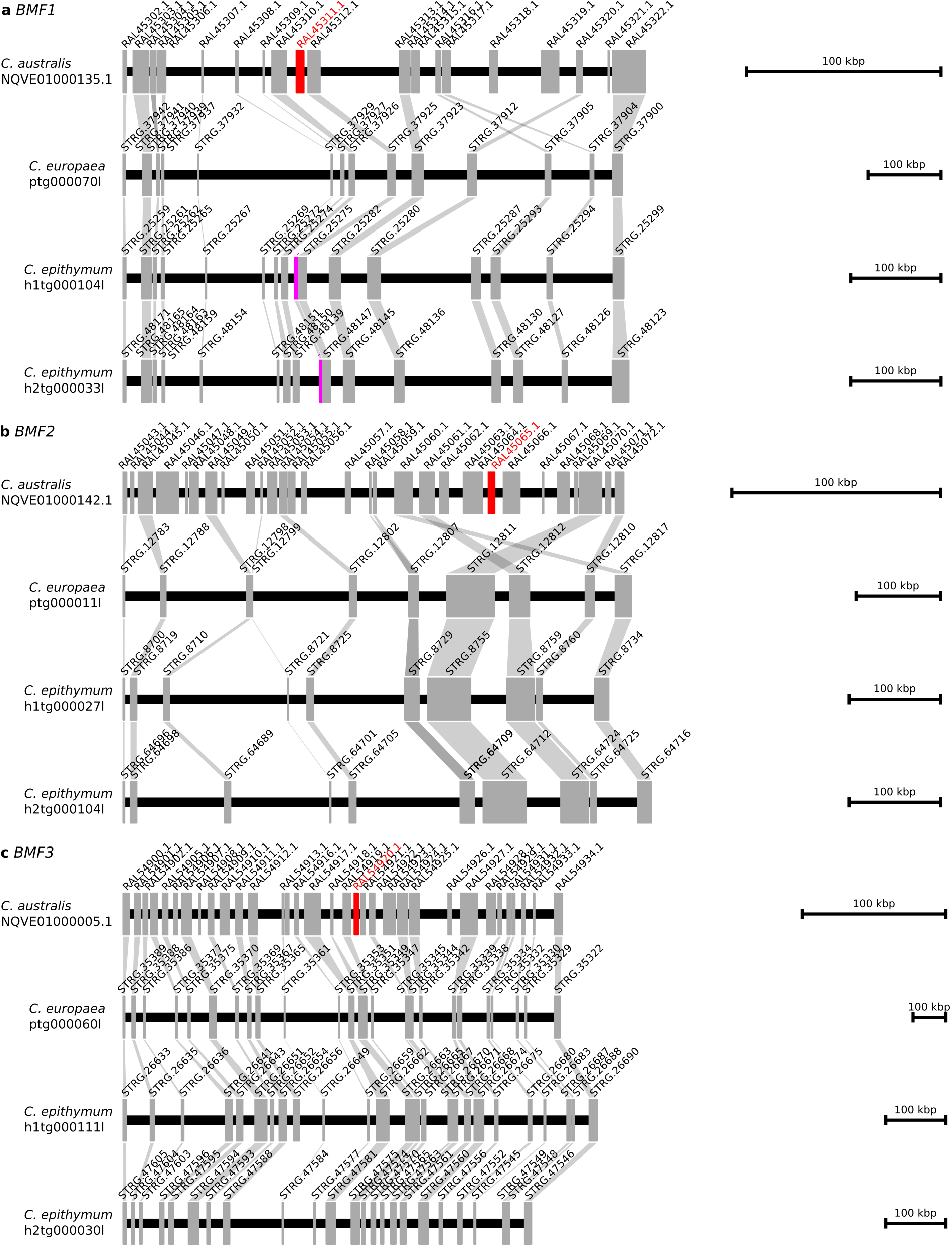
Comparison of orthologous loci, part 2. Comparison of loci possessing *BMF1, BMF2*, and *BMF3* genes in *C. australis* (highlighted in red) with orthologous loci in *C. europaea* and *C. epithymum*. As *C. epithymum* has two haplotypes, each locus is represented by two contigs. Both alleles of *BMF1* gene in *C. epithymum* (highlighted in purple) are truncated and the gene is not transcribed.

**Supplementary Fig. 4.**
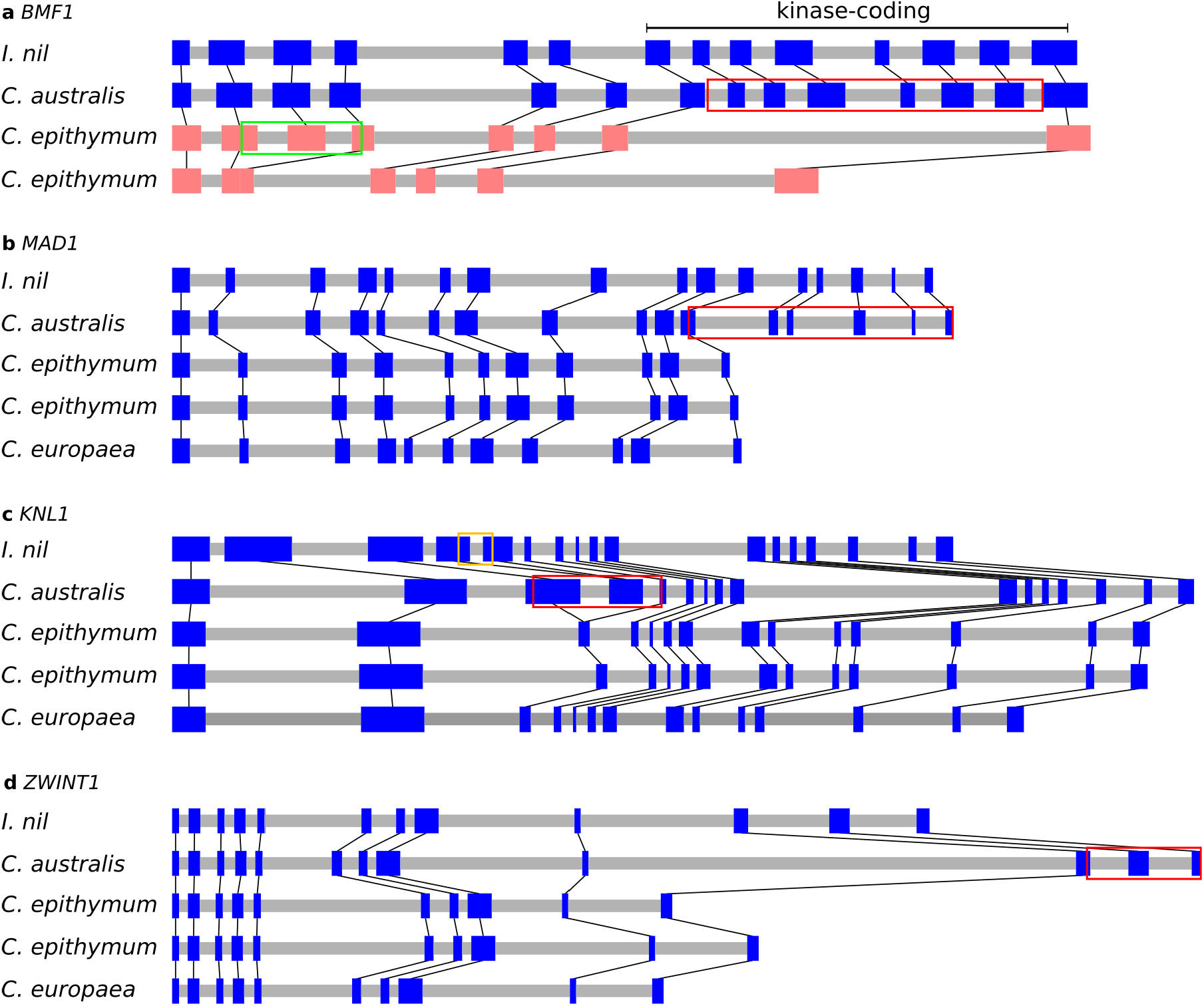
Comparison of exon/intron structures of kinetochore genes that are truncated in holocentric *Cuscuta* species with their full-length homologs in monocentric *C. australis* and *I. nil*. The red rectangles mark exons that are present in *C. australis* but absent or truncated in *C. epithymum*. **a**, Comparison of *BMF1* genes. The exons that were lost in *C. epithymum* encoded kinase domain of BMF1. The green rectangle marks exons present in one allele of the *BMF1* gene that are missing or truncated in the other. As the *BMF1* gene is not transcribed in *C. epithymum*, the exons that remained preserved are not translated into protein. **b-d**, Comparison of *MAD1, KNL1* and *ZWINT1* genes. The orange rectangle in the *I. nil KNL1* gene structure marks a region that is missing in its homolog in *C. australis*.

**Supplementary Fig. 5.**
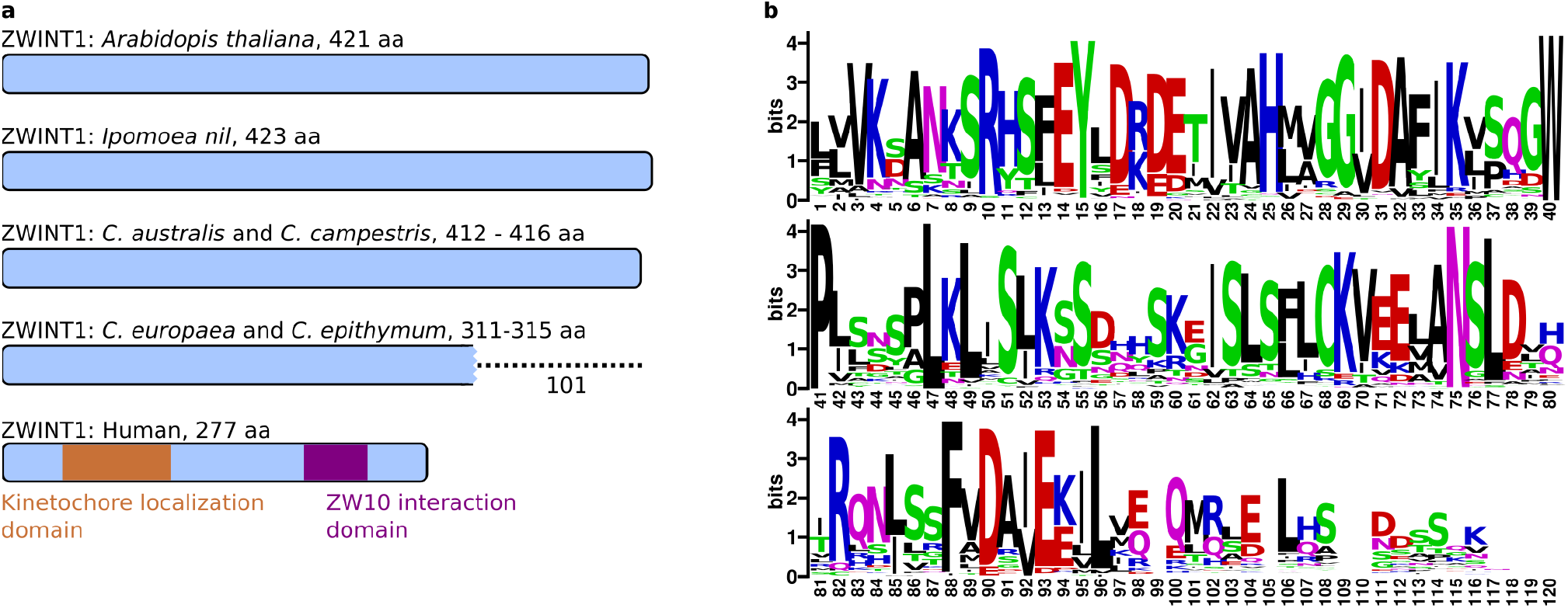
Truncation of ZWINT1 in holocentric *Cuscuta* species. **a**, Schematic of ZWINT1 proteins showing conserved size in monocentric species and C-terminal truncation in *C. europaea* and *C. epithymum* (depicted as a dotted line). As ZWINT1 has not yet been functionally characterized in plants, it is not possible to predict the impact of the truncation. In humans, a domain near the C-terminus interacts with ZW10 protein ^7^, but it shares no sequence similarity with the plant ZWINT1 homologs. **b**, Sequence logo of ZWINT1 C-terminus inferred from alignment of sequences from 129 diverse plant species demonstrating a high level of sequence conservation, suggesting that the ZWINT1 C-terminal domain has a conserved function in plants.

**Supplementary Fig. 6.**
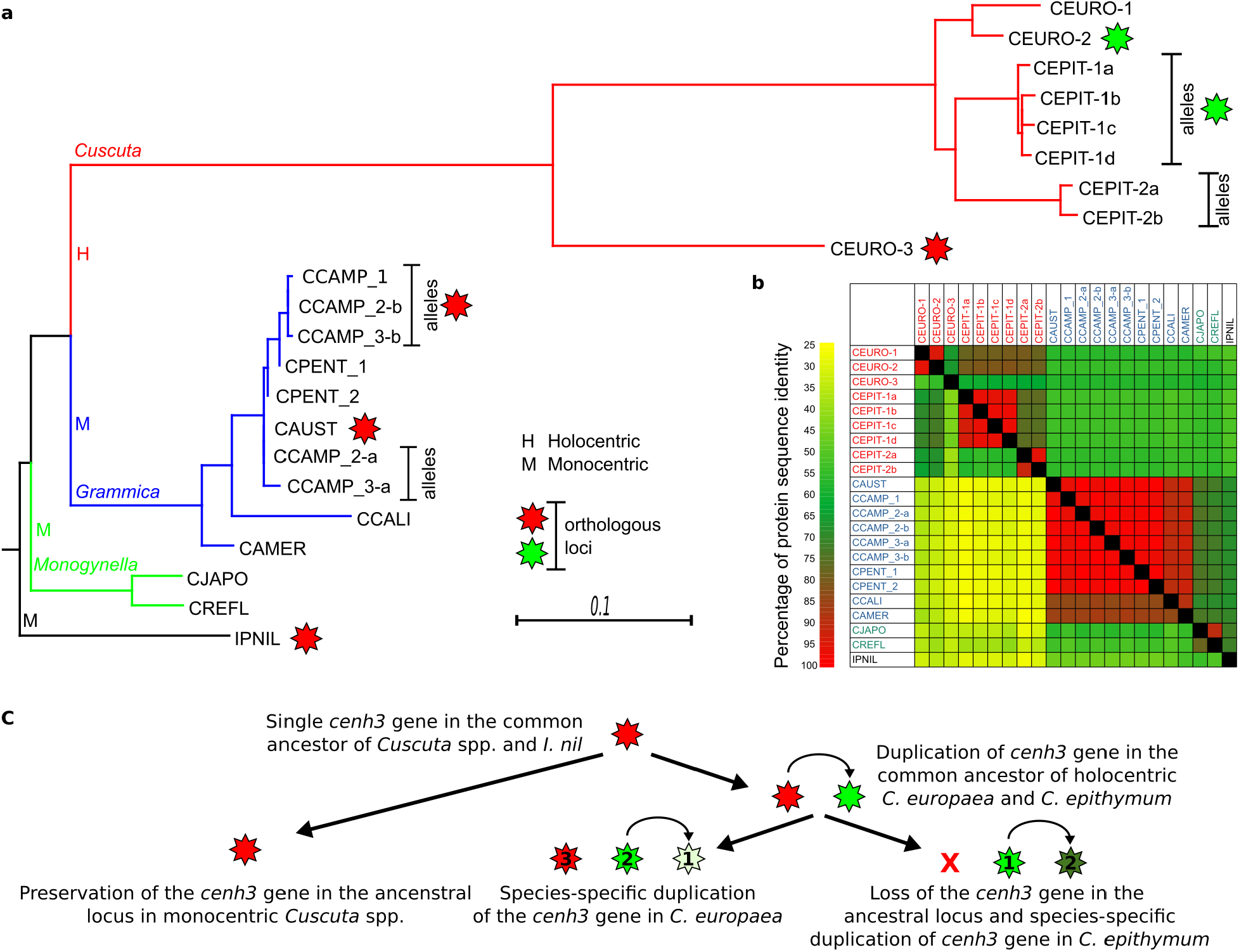
Analysis of CENH3 sequences. **a**, Phylogenetic tree inferred from the alignment of CENH3-coding sequences using the maximum likelihood method, excluding all INDEL sites. Branches corresponding to subgenera *Cuscuta, Grammica*, and *Monogynella*, are colored in red, blue, and green, respectively. Considerably longer branches in the subgenus *Cuscuta* contrast with those in the species trees inferred from *ITS* and *rbcL* sequences (Supplementary Fig. 9) and indicate faster divergence of CENH3 in holocentric compared with monocentric *Cuscuta* species. The sources of the CENH3 sequences used for the analysis are provided in the Supplementary Table 6. **b**, Similarity between CENH3 protein sequences visualized as a heatmap. Boxes above and below the black diagonal show the percentage identity over the entire CENH3 protein sequence and the N-terminus, respectively (the exact values are available in Supplementary Table 9). CENH3 protein sequences in holocentric *Cuscuta* species are considerably more divergent, particularly in the N-terminus, than in monocentric *Cuscuta* species. CENH3 sequences from subgenus *Cuscuta, Grammica*, and *Monogynella* are colored in red, blue, and green, respectively. **c**, Reconstruction of CENH3 gene duplication and loss events in the evolution of holocentric *Cuscuta* species, inferred from the topology of the phylogenetic tree in the panel “a” combined with the information about orthologous CENH3 loci (Supplementary Figs. 7 and 8). These data indicate that the ortholog of *CENH3*^*CEURO-3*^ was lost in *C. epithymum*, and that *CENH3*^*CEURO-1*^ and *CENH3*^*CEPIT-2*^ originated from independent duplications of *CENH3*^*CEURO-2*^ and *CENH3*^*CEPIT-1*^, respectively, which occurred after the divergence of the two species. *CENH3* genes occurring in orthologous loci are indicated by the same star color and the numbers inside the stars indicate the CENH3 variant in respective *Cuscuta* species.

**Supplementary Fig. 7.**
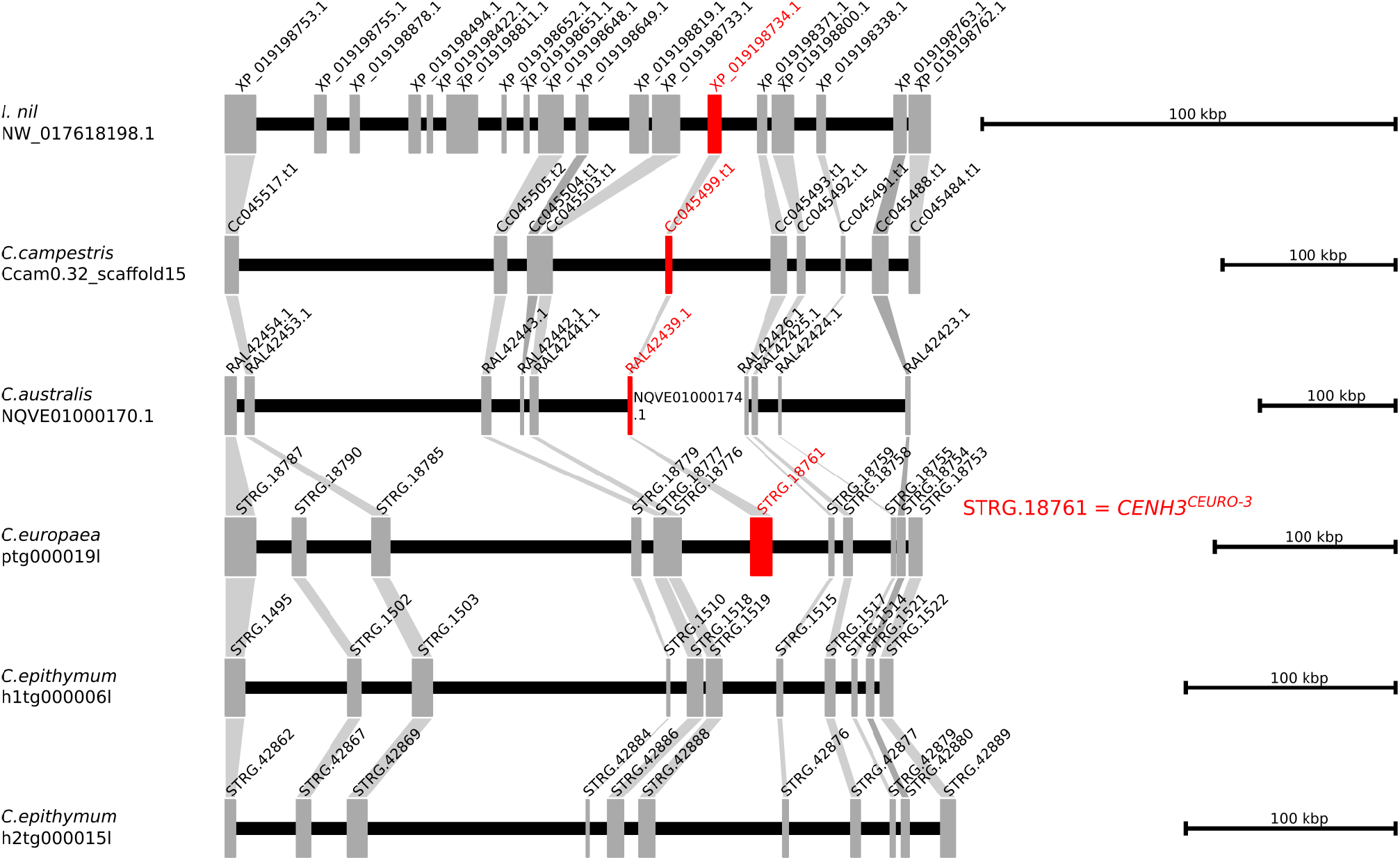
Comparison of *CENH3* gene loci in holocentric *Cuscuta* species. **a**, Comparison of the *CENH3* gene locus in *I. nil* with orthologous loci in *C. australis, C. campestris, C. europaea* and *C. epithymum*. It demonstrates that the all the species except *C. epithymum* maintained the ancestral position of the *CENH3* gene. The *CENH3* genes are highlighted in red.

**Supplementary Fig. 8.**
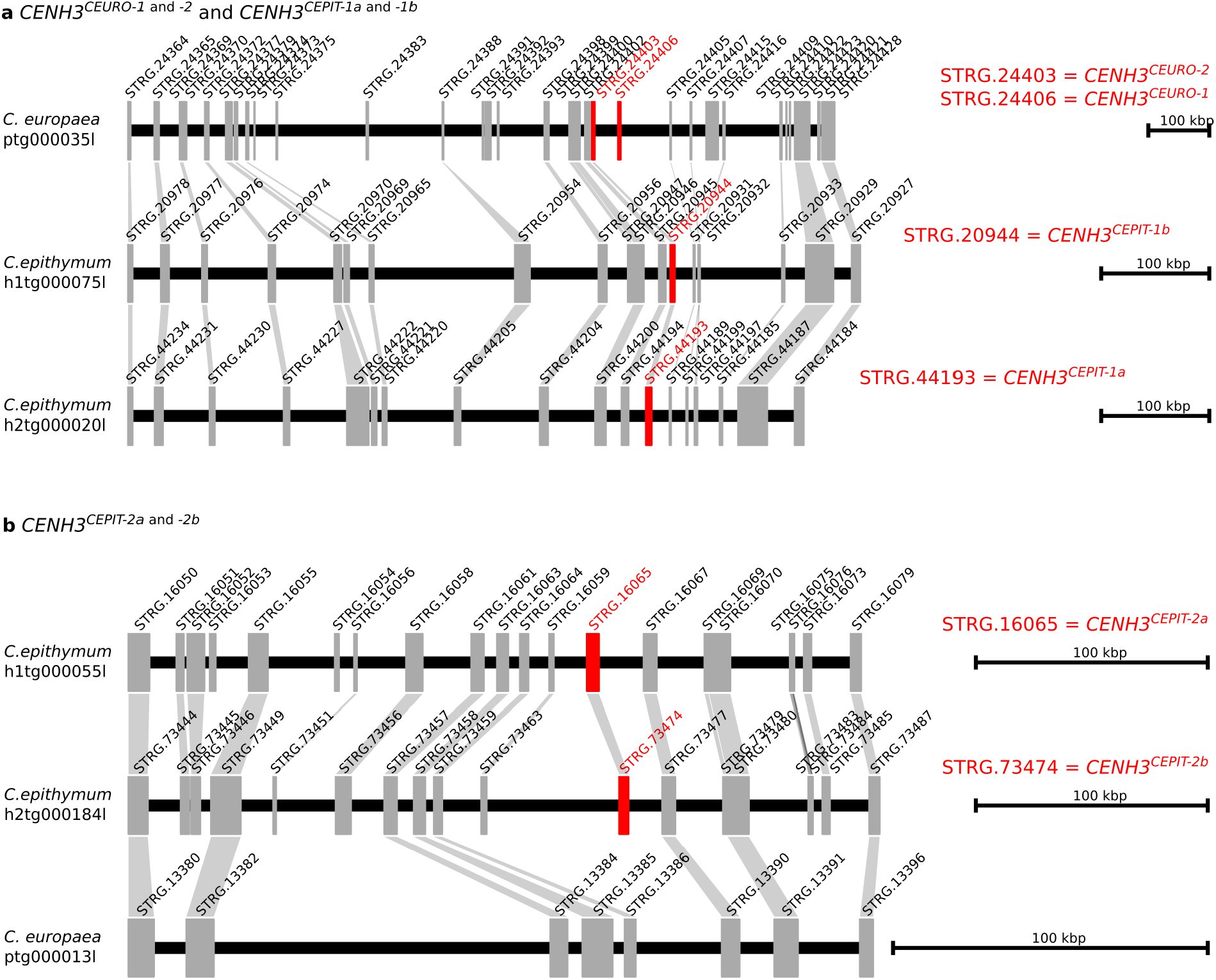
Comparison of CENH3 gene loci between *C. europaea* and *C. epithymum*. **a**, Comparison of the locus containing *CENH3*^*CEURO-1*^ and *CENH3*^*CEURO-2*^ genes in *C. europea* with the orthologous *CENH3*^*CEPIT-1*^ containing locus in *C. epithymum*. **b**, Comparison of the locus containing *CENH3*^*CEPIT-2*^ and the orthologous CENH3-lacking locus in *C. europaea*. The CENH3 genes are highlighted in red.

**Supplementary Fig. 9.**
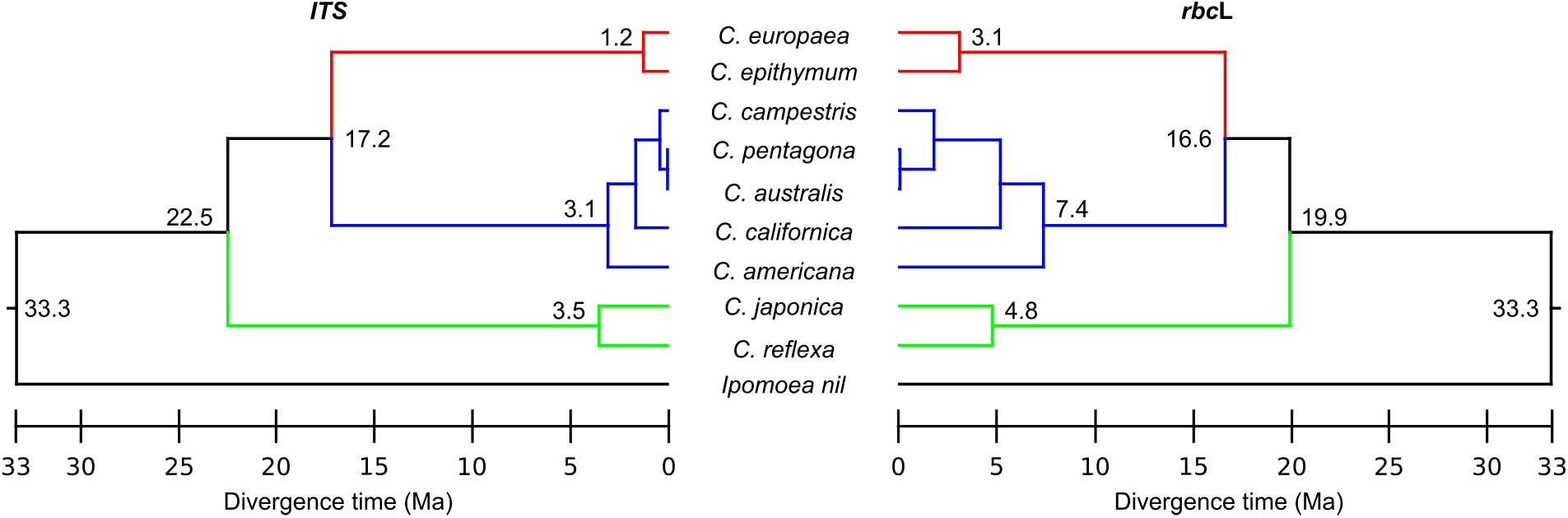
Time-scale phylogenetic trees of *Cuscuta* species included in the study. The trees were inferred from *ITS* (left) and *rbc*L (right) sequences using the maximum likelihood method with smart model selection ^8,9^ and then dated using the RelTime method implemented in MEGA X ^10^, assuming that the most recent common ancestor of *Cuscuta* and *Ipomoea* existed 33.3 million years ago (Ma) ^5^. The numbers at nodes show divergence time in Ma.

**Supplementary Fig. 10.**
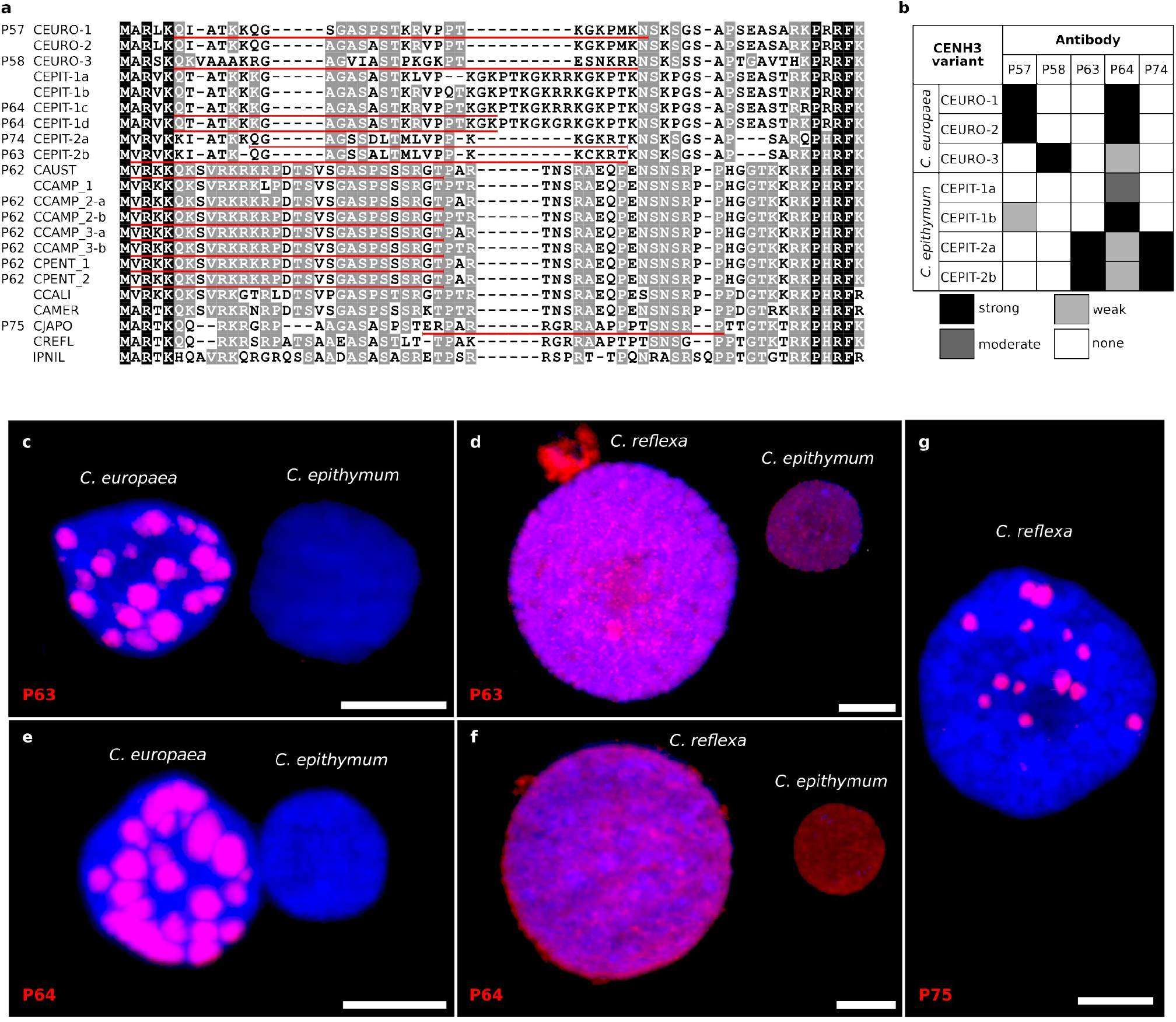
Detection of CENH3 in *C. epithymum*. **a**, Sequence comparison of the peptides used to produce antibodies against CENH3 and the N-terminal sequences of CENH3 variants in *C. europaea* and *C. epithymum*. IDs of the antibodies are shown before the sequence names. **b**, Reactivity of CENH3 antibodies with distinct CENH3 variants in *C. europaea* and *C. epithymum*. The reactivity was tested using western blot detection of CENH3 proteins expressed in *E. coli*. While the different CENH3 antibodies had variable reactivity against individual variants, together they recognized all four CENH3 variants present in the sequenced clone of *C. epithymum*. **c-f**, In situ immunodetection of CENH3. To distinguish specific signals from background, chromosomes and nuclei isolated from *C. epithymum, C. europaea*, and *C. reflexa* were mixed and analyzed on the same slide using the same microscope settings for image acquisition. Because the regions used to generate the antibodies for CENH3 histones from *C. epithymum* showed only partial similarity to CENH3 from *C. europaea* and no significant similarity to CENH3 from *C. reflexa*, the intensity of potential signals in the latter two species could be used to set the threshold for specific signals. The antibodies P63 and P64 strongly labeled the CENH3-containing heterochromatin blocks in *C. europaea* but did not produce a visible signal on chromosomes and nuclei in *C. epithymum* (c,e) at the same exposure time. By contrast, when the exposure time was increased to visualize signals in *C. epithymum*, the signals also appeared on whole nuclei and chromosomes in *C. reflexa* (d,f), indicating that they were not CENH3-specific. The antibody P74 produced no signal in *C. epithymum* and did not label CENH3-containing heterochromatin in *C. europaea* (data not shown). **f**, CENH3 antibody raised to *Monogynella* species (P75) labeled only centromeres in *C. reflexa*. These results suggest that either the amount of CENH3 in chromosomes and nuclei was below the detection limit or that CENH3 was not present in chromatin in *C. epithymum*.

**Supplementary Fig. 11.**
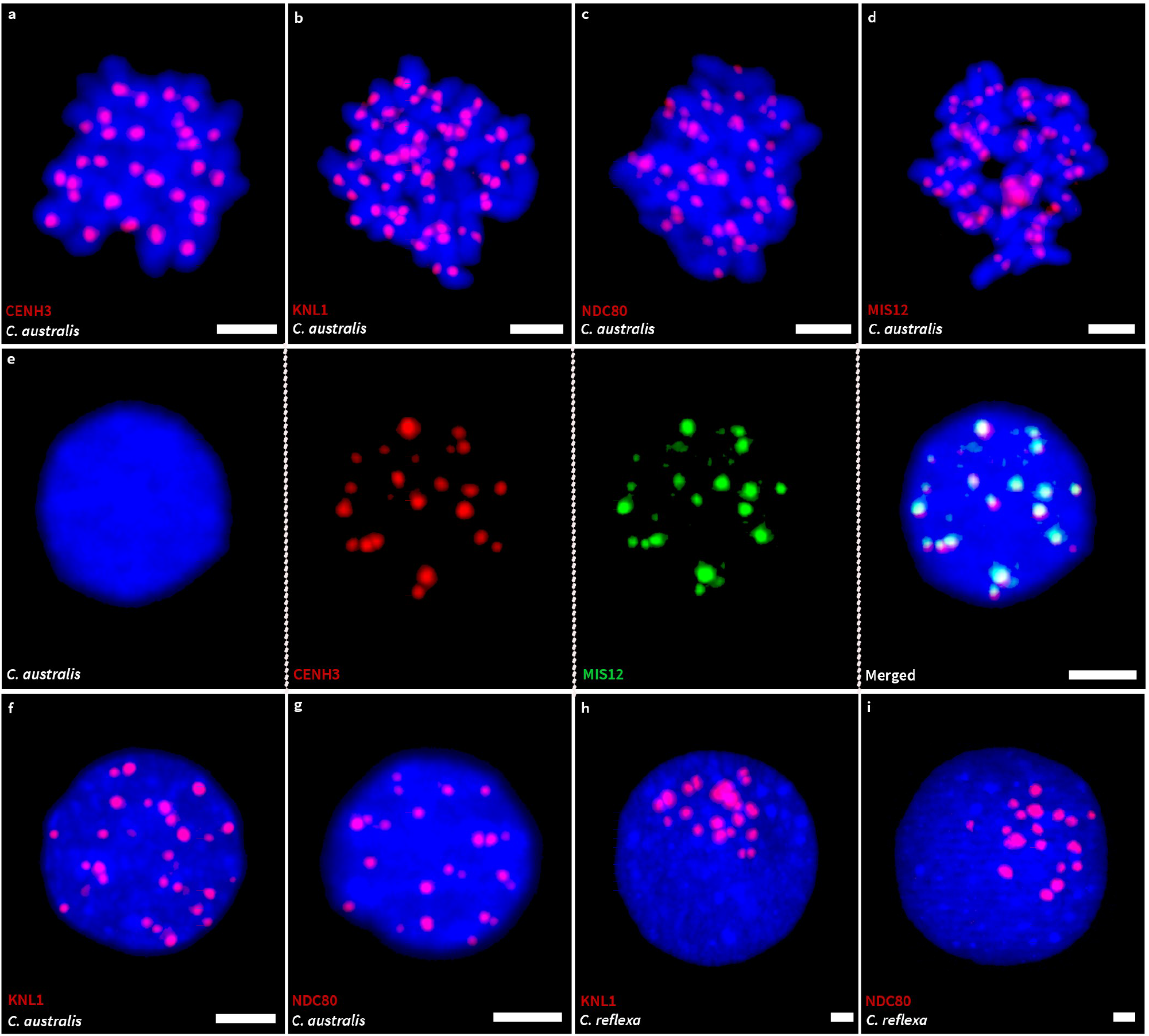
Detection of CENH3 and KMN proteins in *C. australis* and *C. reflexa*. The two species were selected as representatives of monocentric *Cuscuta* species of the subgenus *Grammica* and *Monogynella*, respectively. **a-g**, Detection of kinetochore proteins on mitotic chromosomes (a-d) and nuclei (e-g) in *C. australis*. All kinetochore proteins were detected in a single domain on each sister chromatid of mitotic chromosomes and in discrete centromere domains in interphase nuclei, suggesting that the kinetochore is assembled in centromere domains during all or most of the cell cycle. Simultaneous detection of CENH3 and MIS12 (e) revealed colocalization of the two proteins, confirming the specificity of the MIS12 antibody. **h-i**, Detection of KNL1 and NDC80 in interphase nuclei of *C. reflexa*. The staining pattern resembles that of *C. australis*, suggesting that kinetochores are assembled during interphase in both *Grammica* and *Monogynella* species.

**Supplementary Fig. 12.**
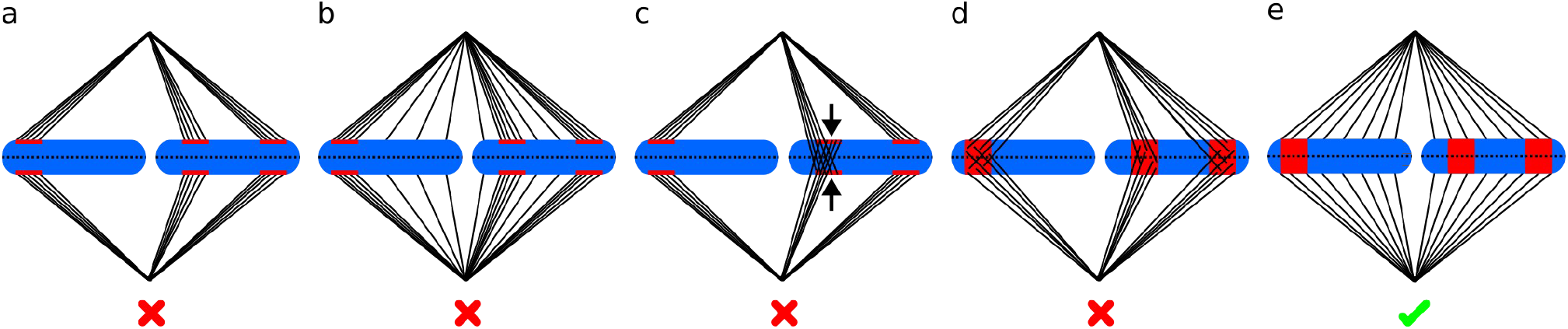
Models for the distribution of CENH3 and tubulin during mitotic metaphase in *C. europaea*. **a-d**, Hypothetical models that would be applicable if CENH3 retained its function as a foundational kinetochore protein, i.e., initiation of kinetochore assembly. Red crosses indicate the models that are not supported by cytogenetic observations. **a**, CENH3 is restricted to the poleward site where microtubules specifically attach to chromosomes. **b**, As for a, but microtubules also attach to chromosomes at sites where CENH3 is present but undetectable. In this case, the density of microtubules would likely be sparse compared with the major sites of CENH3 accumulation. **c**, As for a, except that the presence of two CENH3-containing domains on the same chromatid results in merotelic attachment (arrows). In merotelic attachment, which often occurs in dicentric chromosomes, a single chromatid is attached to microtubules originating from opposite poles. This leads to defects in chromosome segregation. **d**, The presence of CENH3 in transverse bands leads to disordered attachment of chromosomes to mitotic spindle microtubules, which impairs bi-orientation of chromosomes to the mitotic spindle and leads to defects in chromosome segregation. **e**, The observed distribution of CENH3 and microtubules shows that there is no correlation between the density of microtubules of the mitotic spindle and the occurrence of CENH3, suggesting that CENH3 is not a foundational kinetochore protein in mitosis in *C. europaea*.

### Supplementary Movies

**Supplementary Movie 1** | Spatial distribution of CENH3 (red) and KNL1 (green) in an interphase nucleus (blue) of *C. europaea*.

**Supplementary Movie 2** | Spatial distribution of KNL1 (green) and CENP-C (red) in an interphase nucleus (blue) of *C. europaea*.

**Supplementary Movie 3** | Spatial distribution of KNL1 (green) and NDC80 (red) in an interphase nucleus (blue) of *C. europaea*.

**Supplementary Movie 4** | Spatial distribution of CENH3 (red) and MIS12 (green) in an interphase nucleus (blue) of *C. europaea*.

## References

1. Melters, D. P., Paliulis, L. V, Korf, I. F. & Chan, S. W. L. Holocentric chromosomes: convergent evolution, meiotic adaptations, and genomic analysis. Chromosom. Res. 20, 579–93 (2012).

2. McKinley, K. L. & Cheeseman, I. M. The molecular basis for centromere identity and function. Nat. Rev. Mol. Cell Biol. 17, 16–29 (2016).

3. Pesenti, M. E., Weir, J. R. & Musacchio, A. Progress in the structural and functional characterization of kinetochores. Curr. Opin. Struct. Biol. 37, 152–163 (2016).

4. Yamagishi, Y., Sakuno, T., Goto, Y. & Watanabe, Y. Kinetochore composition and its function: lessons from yeasts. FEMS Microbiol. Rev. 38, 185–200 (2014).

5. Lara-Gonzalez, P., Westhorpe, F. G. & Taylor, S. S. The spindle assembly checkpoint. Curr. Biol. 22, R966–R980 (2012).

6. Musacchio, A. The molecular biology of spindle assembly checkpoint signaling dynamics. Curr. Biol. 25, R1002–R1018 (2015).

7. Komaki, S. et al. Functional analysis of the plant chromosomal passenger complex. Plant Physiol. 183, 1586–1599 (2020).

8. Carmena, M., Wheelock, M., Funabiki, H. & Earnshaw, W. C. The chromosomal passenger complex (CPC): from easy rider to the godfather of mitosis. Nat. Rev. Mol. Cell Biol. 13, 789–803 (2012).

9. van der Waal, M. S., Hengeveld, R. C. C., van der Horst, A. & Lens, S. M. A. Cell division control by the Chromosomal Passenger Complex. Exp. Cell Res. 318, 1407–1420 (2012).

10. Buchwitz, B. J., Ahmad, K., Moore, L. L., Roth, M. B. & Henikoff, S. A histone-H3-like protein in C. elegans. Nature 401, 547–548 (1999).

11. Schubert, V. et al. Super-resolution microscopy reveals diversity of plant centromere architecture. Int. J. Mol. Sci. 21, 3488 (2020).

12. Cortes-Silva, N. et al. CenH3-independent kinetochore assembly in Lepidoptera requires CCAN, including CENP-T. Curr. Biol. 30, 561-572.e10 (2020).

13. Drinnenberg, I. A., DeYoung, D., Henikoff, S. & Malik, H. S. Recurrent loss of CenH3 is associated with independent transitions to holocentricity in insects. Elife 3, e03676 (2014).

14. Senaratne, A. P. et al. Formation of the CenH3-deficient holocentromere in Lepidoptera avoids active chromatin. Curr. Biol. 31, 173-181.e7 (2021).

15. Oliveira, L. et al. Mitotic spindle attachment to the holocentric chromosomes of Cuscuta europaea does not correlate with the distribution of CENH3 chromatin. Front. Plant Sci. 10, 1799 (2020).

16. Neumann, P. et al. Impact of parasitic lifestyle and different types of centromere organization on chromosome and genome evolution in the plant genus Cuscuta. New Phytol. 229, 2365–2377 (2021).

17. Zuo, S. et al. Recurrent plant-specific duplications of KNL2 and its conserved function as a kinetochore assembly factor. Mol. Biol. Evol. 39, (2022).

18. Maddox, P. S., Hyndman, F., Monen, J., Oegema, K. & Desai, A. Functional genomics identifies a Myb domain-containing protein family required for assembly of CENP-A chromatin. J. Cell Biol. 176, 757–763 (2007).

19. Steiner, F. a & Henikoff, S. Holocentromeres are dispersed point centromeres localized at transcription factor hotspots. Elife 3, 1–22 (2014).

20. Lermontova, I. et al. Arabidopsis KINETOCHORE NULL2 is an upstream component for centromeric Histone H3 Variant cenH3 deposition at centromeres. Plant Cell 25, 3389–404 (2013).

21. Pintard, L. & Bowerman, B. Mitotic cell division in Caenorhabditis elegans. Genetics 211, 35–73 (2019).

22. Vondrak, T. et al. Complex sequence organization of heterochromatin in the holocentric plant Cuscuta europaea elucidated by the computational analysis of nanopore reads. Comput. Struct. Biotechnol. J. 19, 2179–2189 (2021).

23. Dimitrova, Y. N., Jenni, S., Valverde, R., Khin, Y. & Harrison, S. C. Structure of the MIND complex defines a regulatory focus for yeast kinetochore assembly. Cell 167, 1014-1027.e12 (2016).

24. Petrovic, A. et al. Structure of the MIS12 complex and molecular basis of its interaction with CENP-C at human kinetochores. Cell 167, 1028-1040.e15 (2016).

25. Screpanti, E. et al. Direct binding of Cenp-C to the Mis12 complex joins the inner and outer kinetochore. Curr. Biol. 21, 391–398 (2011).

26. Przewloka, M. R. et al. CENP-C is a structural platform for kinetochore assembly. Curr. Biol. 21, 399–405 (2011).

27. Tromer, E. C., Wemyss, T. A., Ludzia, P., Waller, R. F. & Akiyoshi, B. Repurposing of synaptonemal complex proteins for kinetochores in Kinetoplastida. Open Biol. 11, (2021).

28. Butenko, A. et al. Evolution of metabolic capabilities and molecular features of diplonemids, kinetoplastids, and euglenids. BMC Biol. 18, 1–28 (2020).

29. Karg, T., Elting, M. W., Vicars, H., Dumont, S. & Sullivan, W. The chromokinesin Klp3a and microtubules facilitate acentric chromosome segregation. J. Cell Biol. 216, 1597–1608 (2017).

30. Vicars, H., Karg, T., Warecki, B., Bast, I. & Sullivan, W. Kinetochore-independent mechanisms of sister chromosome separation. PLOS Genet. 17, e1009304 (2021).

31. Swentowsky, K. W. et al. Distinct kinesin motors drive two types of maize neocentromeres. Genes Dev. 34, 1239–1251 (2020).

32. Dawe, R. K. et al. A Kinesin-14 motor activates neocentromeres to promote meiotic drive in maize. Cell 173, 839–850 (2018).

33. Dellaporta, S. L. S. L., Wood, J. & Hicks, J. B. J. B. A plant DNA minipreparation: version II. Plant Mol. Biol. Report. 1, 19–21 (1983).

34. Vondrak, T. et al. Characterization of repeat arrays in ultra-long nanopore reads reveals frequent origin of satellite DNA from retrotransposon-derived tandem repeats. Plant J. 101, 484–500 (2020).

35. Zimin, A. V. et al. The MaSuRCA genome assembler. Bioinformatics 29, 2669–2677 (2013).

36. Cheng, H., Concepcion, G. T., Feng, X., Zhang, H. & Li, H. Haplotype-resolved de novo assembly using phased assembly graphs with hifiasm. Nat. Methods 18, 170–175 (2021).

37. Manni, M., Berkeley, M. R., Seppey, M., Simão, F. A. & Zdobnov, E. M. BUSCO update: novel and streamlined workflows along with broader and deeper phylogenetic coverage for scoring of eukaryotic, prokaryotic, and viral genomes. Mol. Biol. Evol. 38, 4647–4654 (2021).

38. Gurevich, A., Saveliev, V., Vyahhi, N. & Tesler, G. QUAST: quality assessment tool for genome assemblies. Bioinformatics 29, 1072–1075 (2013).

39. Marçais, G. & Kingsford, C. A fast, lock-free approach for efficient parallel counting of occurrences of k-mers. Bioinformatics 27, 764–770 (2011).

40. Vurture, G. W. et al. GenomeScope: fast reference-free genome profiling from short reads. Bioinformatics 33, 2202–2204 (2017).

41. Grabherr, M. G. et al. Full-length transcriptome assembly from RNA-Seq data without a reference genome. Nat. Biotechnol. 29, 644–652 (2011).

42. Dobin, A. et al. STAR: ultrafast universal RNA-seq aligner. Bioinformatics 29, 15–21 (2013).

43. Li, H. et al. The sequence alignment/map format and SAMtools. Bioinformatics 25, 2078–2079 (2009).

44. Pertea, M. et al. StringTie enables improved reconstruction of a transcriptome from RNA-seq reads. Nat. Biotechnol. 33, 290–295 (2015).

45. Emms, D. M. & Kelly, S. OrthoFinder: phylogenetic orthology inference for comparative genomics. Genome Biol. 20, 1–14 (2019).

46. Dunn, N. A. et al. Apollo: democratizing genome annotation. PLOS Comput. Biol. 15, e1006790 (2019).

47. Komaki, S. & Schnittger, A. The spindle assembly checkpoint in Arabidopsis is rapidly shut off during severe stress. Dev. Cell 43, 172-185.e5 (2017).

48. Su, H. et al. Knl1 participates in spindle assembly checkpoint signaling in maize. Proc. Natl. Acad. Sci. 118, e2022357118 (2021).

49. van Hooff, J. J., Tromer, E., van Wijk, L. M., Snel, B. & Kops, G. J. Evolutionary dynamics of the kinetochore network in eukaryotes as revealed by comparative genomics. EMBO Rep. 18, 1559–1571 (2017).

50. Sun, G. et al. Large-scale gene losses underlie the genome evolution of parasitic plant Cuscuta australis. Nat. Commun. 9, 2683 (2018).

51. Vogel, A. et al. Footprints of parasitism in the genome of the parasitic flowering plant Cuscuta campestris. Nat. Commun. 9, 2515 (2018).

52. Hoshino, A. et al. Genome sequence and analysis of the Japanese morning glory Ipomoea nil. Nat. Commun. 7, 13295 (2016).

53. Brankovics, B. et al. GRAbB: selective assembly of genomic regions, a new niche for genomic research. PLOS Comput. Biol. 12, e1004753 (2016).

54. Birney, E., Clamp, M. & Durbin, R. GeneWise and Genomewise. Genome Res. 14, 988–995 (2004).

55. Edgar, R. C. MUSCLE: multiple sequence alignment with high accuracy and high throughput. Nucleic Acids Res. 32, 1792–1797 (2004).

56. Novák, P., Neumann, P. & Macas, J. Global analysis of repetitive DNA from unassembled sequence reads using RepeatExplorer2. Nat. Protoc. 15, 3745–3776 (2020).

57. Bailey, T. L. & Elkan, C. Fitting a mixture model by expectation maximization to discover motifs in biopolymers. Proceedings. Int. Conf. Intell. Syst. Mol. Biol. 2, 28–36 (1994).

58. Crooks, G. E., Hon, G., Chandonia, J.-M. & Brenner, S. E. WebLogo: a sequence logo generator. Genome Res. 14, 1188–90 (2004).

59. Weisshart, K., Fuchs, J. & Schubert, V. Structured illumination microscopy (SIM) and photoactivated localization microscopy (PALM) to analyze the abundance and distribution of RNA polymerase II molecules on flow-sorted Arabidopsis nuclei. Bio-protocol 6, e1725 (2016).

60. Hara, M. & Fukagawa, T. Where is the right path heading from the centromere to spindle microtubules? Cell Cycle 18, 1199–1211 (2019).

61. Ciferri, C. et al. Implications for kinetochore-microtubule attachment from the structure of an engineered Ndc80 Complex. Cell 133, 427–439 (2008).

62. Alushin, G. M. et al. The Ndc80 kinetochore complex forms oligomeric arrays along microtubules. Nature 467, 805–810 (2010).

63. Welburn, J. P. I. et al. Aurora B phosphorylates spatially distinct targets to differentially regulate the kinetochore-microtubule interface. Mol. Cell 38, 383–392 (2010).

64. Petrovic, A. et al. Modular assembly of RWD domains on the Mis12 complex underlies outer kinetochore organization. Mol. Cell 53, 591–605 (2014).

65. Valverde, R., Ingram, J. & Harrison, S. C. Conserved tetramer junction in the kinetochore Ndc80 complex. Cell Rep. 17, 1915–1922 (2016).

66. Ali‐Ahmad, A., Bilokapić, S., Schäfer, I. B., Halić, M. & Sekulić, N. CENP-C unwraps the human CENP-A nucleosome through the H2A C-terminal tail. EMBO Rep. 20, 1–13 (2019).

67. Hornung, P. et al. A cooperative mechanism drives budding yeast kinetochore assembly downstream of CENP-A. J. Cell Biol. 206, 509–524 (2014).

68. Ghongane, P., Kapanidou, M., Asghar, A., Elowe, S. & Bolanos-Garcia, V. M. The dynamic protein Knl1 - a kinetochore rendezvous. J. Cell Sci. 127, 3415–3423 (2014).

69. Caillaud, M. C. et al. Spindle assembly checkpoint protein dynamics reveal conserved and unsuspected roles in plant cell division. PLoS One 4, (2009).

70. Zhang, H. et al. Role of the BUB3 protein in phragmoplast microtubule reorganization during cytokinesis. Nat. Plants 4, 485–494 (2018).

71. Luo, Y., Ahmad, E. & Liu, S.-T. MAD1: kinetochore receptors and catalytic mechanisms. Front. Cell Dev. Biol. 6, 1–10 (2018).

## Supplementary References

1. Zimin, A. V. et al. The MaSuRCA genome assembler. Bioinformatics 29, 2669–2677 (2013).

2. Cheng, H., Concepcion, G. T., Feng, X., Zhang, H. & Li, H. Haplotype-resolved de novo assembly using phased assembly graphs with hifiasm. Nat. Methods 18, 170–175 (2021).

3. Neumann, P. et al. Impact of parasitic lifestyle and different types of centromere organization on chromosome and genome evolution in the plant genus Cuscuta. New Phytol. 229, 2365–2377 (2021).

4. Howley, P. M., Israel, M. a, Law, M. F. & Martin, M. a. A rapid method for detecting and mapping homology between heterologous DNAs. Evaluation of polyomavirus genomes. J. Biol. Chem. 254, 4876–83 (1979).

5. Sun, G. et al. Large-scale gene losses underlie the genome evolution of parasitic plant Cuscuta australis. Nat. Commun. 9, 2683 (2018).

6. Vogel, A. et al. Footprints of parasitism in the genome of the parasitic flowering plant Cuscuta campestris. Nat. Commun. 9, 2515 (2018).

7. Vos, L. J., Famulski, J. K. & Chan, G. K. T. HZwint-1 bridges the inner and outer kinetochore: identification of the kinetochore localization domain and the hZw10-interaction domain. Biochem. J. 436, 157–168 (2011).

8. Lefort, V., Longueville, J. E. & Gascuel, O. SMS: smart model selection in PhyML. Mol. Biol. Evol. 34, 2422–2424 (2017).

9. Guindon, S. et al. New algorithms and methods to estimate maximum-likelihood phylogenies: assessing the performance of PhyML 3.0. Syst. Biol. 59, 307–321 (2010).

10. Mello, B. Estimating TimeTrees with MEGA and the TimeTree resource. Mol. Biol. Evol. 35, 2334–2342 (2018).

